# Reconstitution of Human Brain Cell Diversity in Organoids via Four Protocols

**DOI:** 10.1101/2024.11.15.623576

**Authors:** Julia Naas, Meritxell Balmãna, Laurenz Holcik, Maria Novatchkova, Lina Dobnikar, Thomas Krausgruber, Sabrina Ladstätter, Christoph Bock, Arndt von Haeseler, Christopher Esk, Jürgen A. Knoblich

**Affiliations:** Center for Integrative Bioinformatics Vienna (CIBIV), Max Perutz Labs, University of Vienna and Medical University of Vienna, Vienna BioCenter, Vienna, Austria; Vienna BioCenter PhD Program, a Doctoral School of the University of Vienna and Medical University of Vienna, Vienna, Austria; Institute of Molecular Biotechnology of the Austrian Academy of Science (IMBA), Vienna BioCenter (VBC), Vienna, Austria; Institute of Molecular Pathology (IMP), Vienna, Austria; CeMM Research Center for Molecular Medicine of the Austrian Academy of Sciences, Vienna, Austria; Medical University of Vienna, Institute of Artificial Intelligence, Center for Medical Data Science, Vienna, Austria; Bioinformatics and Computational Biology, Faculty of Computer Science, University of Vienna, Vienna, Austria; Ludwig Boltzmann Institute for Network Medicine at the University of Vienna, Vienna, Austria; Institute of Molecular Biology, University of Innsbruck, Innsbruck, Austria; Medical University of Vienna, Vienna BioCenter (VBC), Vienna, Austria

**Keywords:** brain development, brain organoid, scRNA-seq, protocols, computational tool, data explorer

## Abstract

Human brain organoids are powerful *in vitro* models for brain development and disease. However, their variability can complicate use in biomedical research and drug discovery. Both the specific protocol as well as the pluripotent starting cell line influence organoid variability and can result in incomplete representation of brain cell types in an organoid experiment. Here, we systematically analyze the cellular and transcriptional landscape of brain organoids grown from multiple cell lines using four different protocols recapitulating dorsal and ventral forebrain, midbrain, and striatum. We establish the NEST-Score as a quantitative readout for cell line-driven and protocol-driven differentiation propensities by comparing cellular states across multiple cell lines and to *in vivo* reference data sets. Thereby, we establish a set of organoid protocols that together recreate the vast majority of cell types in the developing human brain and provide a reference for how well cell types are recapitulated across cell lines in each protocol. Additionally, we survey factors contributing to variability during organoid development and identify early gene expression signatures predicting protocol-driven organoid generation at later stages. We provide easy online access to our data through a web-based analysis tool, creating a reference for brain organoid research that allows rapid, straightforward validation of protocol and cell line performance.

## Main Text

Human brain organoids are three-dimensional, self-organizing *in vitro* models that recapitulate functional as well as structural aspects of the developing brain (1–13). In order to capture the neuronal complexity of the brain, a variety of protocols have been established to recapitulate the development of specific brain regions, such as dorsal or ventral forebrain or midbrain (14–19). Publications describing organoid protocols typically include evidence showing that the right cell types are generated, but variability across samples and cell lines has been a challenge (13, 20–22). Cell lines can have intrinsic biases so that individual cell types are created in variable amounts. Furthermore, a strong cell line bias can also lead to the formation of tissue consisting of cell types that are not present in the tissue to be modeled. Without any additional information, it is impossible to determine whether a transcriptomic state of an organoid’s cell is driven by cell line intrinsic properties or the guidance cues of a protocol. This poses a challenge to any study using patient-derived cell lines as it is unclear whether phenotypic effects are consequences of the individual genetic background, disease-associated genetic variants or due to other cell line or protocol driven variability. In an effort to define reference protocol signatures that are independent of the specific cell line used, we systematically evaluated four organoid growth protocols based on two success factors: (1) The reliable generation of the same cell types across multiple cell lines and (2) the generation of cell types that form the respective target brain region *in vivo*. Towards this goal, we provide a comprehensive dataset that evaluates (1) intrinsic biases of cell lines within a given brain organoid protocol, (2) reliability of protocols across multiple cell lines and (3) similarity between the cells generated *in vitro* and *in vivo*. We examined four protocols guiding organoid growth towards dorsal forebrain, ventral forebrain, midbrain, and striatum by conducting single-cell RNA sequencing (scRNA-seq) experiments on mature organoids and time-resolved bulk RNA-sequencing. For each protocol, we distinguish protocol-driven cell states formed across multiple cell lines from cell line-driven cell states that are only observed in few cell lines. Thereby, we can determine the reliable generation of cell types, the first key success factor of a protocol. In addition, we measure how well brain organoid cell types match *in vivo* counterparts as second success factor. To quantify both success factors we have developed a scoring system, the NEST-Score (NEighbourhood Sample homogeneity-Score), that evaluates how well different samples cover the transcriptomic state of each cell based on its neighborhood in Principal Component space. In each protocol, we demonstrate that more than half of the considered cell lines contribute predominantly to protocol-driven cell states, the vast majority of which recapitulates cells found *in vivo* (23, 24). To understand, whether the success of a protocol can already be accessed at early stages, we performed time-resolved bulk RNAsequencing across all protocols and seven cell lines. This allows us to obtain cell line independent, protocol-specific marker gene sets that may serve as a reference for future brain organoid experiments. We provide easy open access to our data via a web-based data explorer, which enables simultaneous browsing across both scRNA and bulk RNA-seq data for individual genes or gene groups of interest (https://vienna-brain-organoid-explorer.vbc.ac.at).

### A brain organoid single-cell RNA-sequencing dataset across protocols and cell lines

To explore a large variety of cell types generated *in vitro*, we employed four distinct brain organoid protocols designed to generate dorsal and ventral forebrain, midbrain and striatum tissue using eight different cell lines for organoid generation. We examined the cellular composition of 120-day old organoids in duplicates using droplet-based scRNA-sequencing (10X Genomics) (Fig. 1A, Fig. S1A). Organoids were grown to day 120 because at that time point, most major mature neuronal cell types are created while a diverse population of progenitor cells is retained (13, 27–29). Cell types present in various brain organoids included deep and upper layer excitatory neurons, different subtypes of inhibitory neurons, as well as glia and oligodendrocyte precursors. All conditions were grown and sequenced in parallel to minimize batch variability, eliminating the need for subsequent computational batch correction. Quality control filtering (30) retained approximately 70 000 high-quality single cell transcriptomes for downstream analysis (see Methods) (Fig. 1B, Fig. S1A). High-quality cells in the combined dataset encompassing all protocols and cell lines experiments reveal cell clusters that were annotated for cell types and brain region based on marker gene expression and previous literature (13, 20, 27, 29) and visualized using uniform manifold approximation and projection (UMAP). We identified a wide range of glial and neuronal cell types in distinct clusters that reflect neuron maturation (Fig. 1B, Sup. Table 1, Sup. Table 2). Developmental trajectories were preserved, and glia and neurons from different dorsal, ventral, midbrain and striatal brain regions clearly distinct. For instance, the well-documented differentiation of excitatory neurons from dorsal radial glia (marked by PAX6, HOPX expression) through intermediate progenitors (EOMES expression) into immature, deep and upper layer neurons (marked by SLA, TBR1 and SATB2 expression, respectively) form a distinct set of clusters. This trajectory is separate from ventral, inhibitory progenitors and neurons (DLX2, SST expression) as well as midbrain-like cells (FOXA2, EN1 expression). Additionally, some cells of non-neural origin such as muscle (smooth muscle marker DES) and stromal cells (fibroblast marker COL3A1) can be observed (Fig. S1B). In the combined UMAP, cells derived from dorsal, ventral and midbrain protocols occupy different transcriptional states (Fig. 1C, Fig. S1C). Cells of the exci-tatory neuronal lineage originated from organoids grown in the dorsal protocol, while the ventral protocol predominantly contributed to the development of inhibitory neurons. The midbrain protocol, on the other hand, generated cells that follow a midbrain-specific neuronal developmental lineage derived from floor plate progenitor cells (Fig. 1B, C, Fig. S1C). In contrast, the striatum protocol produced cell states that were also found in the other three protocols, but additionally generated unique cells not produced by the other three protocols. These striatum-protocol specific cells express markers consistent with medium spiny neuron identity (SIX3, SP9) (Fig. S1B). Overall, the four differentiation protocols across all cell lines collectively yield a highly diverse array of cell types, underscoring the potential of brain organoid protocols to model many neural cell types in the developing human brain (Fig. 1D).

**Fig. 1.**
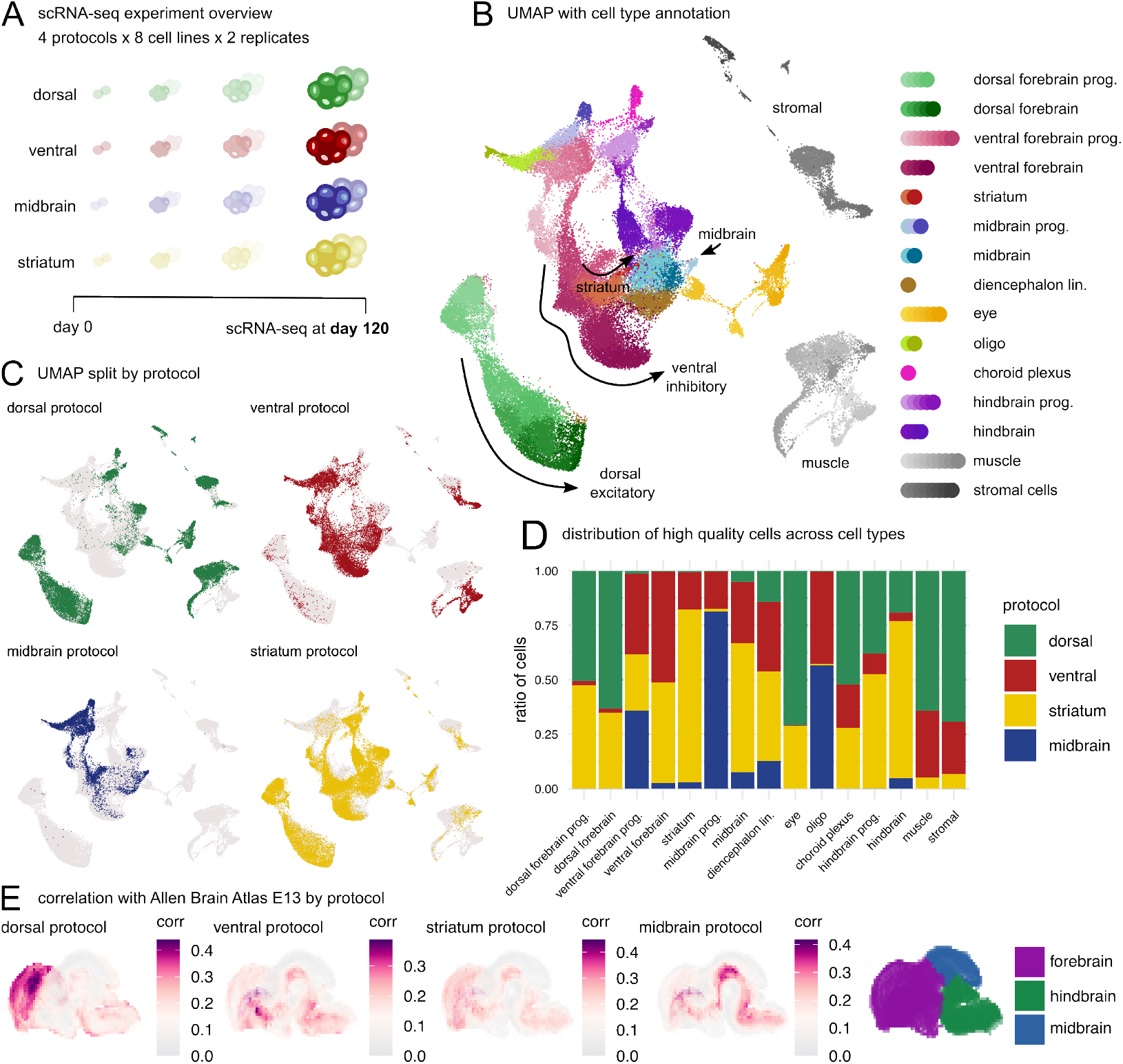
The transcriptional potential of organoids grown from multiple protocols and cell lines at single-cell resolution. (A) Experimental overview for endpoint scRNA-seq analysis of brain organoids from 8 cell lines, 4 protocols. (B) UMAP embedding of scRNA-seq data across four protocols and eight cell lines, color-coded by cell types. Abbreviations: prog. = progenitors, lin. = lineage (C) UMAP of scRNA-seq data colored by the protocol of origin. (D) Contribution of protocols to annotated cell types of the scRNA-seq dataset. (E) VoxHunt (25) analysis of organoid scRNA-seq data with spatial transcriptomics data of a mouse brain slice at embryonic day 13 (BrainSpan, (26)) shows protocol-specific Spearman correlation patterns.

To link the cells produced in the four protocols to their corresponding anatomical locations, we correlated our data with spatially resolved transcriptome data from comparable mouse brain slices (BrainSpan, (26)) using VoxHunt, a tool previously used to assess regional identity in brain organoids (25) (Fig. 1E). Cells from each protocol generally match well to the corresponding anatomical mouse brain region, indicating that the organoids are patterned as expected. However, upon closer inspection, minor positive correlation of gene expression profiles was also observed with unexpected brain regions, such as cells from the midbrain protocol correlating with mouse ventral forebrain structures. Indeed, cell type clusters in the combined UMAP annotated to be midbrain-like correlated better with the corresponding BrainSpan / VoxHunt midbrain reference than the sum of midbrain protocol derived cells, indicating that protocol subsets contained additional non-targeted cells (compare Fig. 1E and Fig. S1D). This observation led us to split the combined UMAP by both individual cell line and protocol combinations (Fig. S1E). This revealed that individual cell lines vary in the degree to which they contribute to cell types generated with each protocol. While most cell lines generated cells consistent with cell types targeted by the respective protocols, some produced aberrant cell types. In the midbrain protocol, for example, cell line Uofv_1 produced a considerable number of inhibitory neurons that align with those generated in the ventral forebrain protocol. In the dorsal protocol, cell lines 176 and Xuja_2 gave rise to different non-neural tissues (muscle and stromal cells). As such unintended cell types were generated only from few cell lines, they can be recognized by a low degree of intermixing with cells from most other cell lines. This indicates that inherent tendencies of certain cell lines can override the guidance provided by a protocol, prompting us to categorize these instances by determining whether a cell state is consistently produced within a protocol.

### Definition of Protocol-Driven and Cell Line-Driven Cell States

To assess how reliable specific cell types are generated in one protocol from multiple cell lines, we introduce the cellular NEST-Score (NEighborhood Sample homogeneiTyScore) (Fig. 2A, Methods). Computed for each individual cell, this score measures the degree to which a cell’s neigh-borhood is composed of cells from different cell lines. A cell’s maximum score of “0” indicates that this cell’s neighborhood is consisting perfectly of all cell lines in the experiment, while a cell’s low (negative) NEST-Score indicates that neighboring cells are derived predominantly from the same cell line. Therefore, high NEST-Scores indicate that corresponding cells are consistently produced within a given protocol as multiple cell lines generate similar cell states. Vice versa, a cell’s low NEST-Score indicates a cell-line driven, non-protocol-conform cell state mainly generated from a particular cell line. It is important to note that this assessment is independent of how well a cell corresponds to the targeted *in vivo* counterpart (see below for *in vivo* comparisons).

**Fig. 2.**
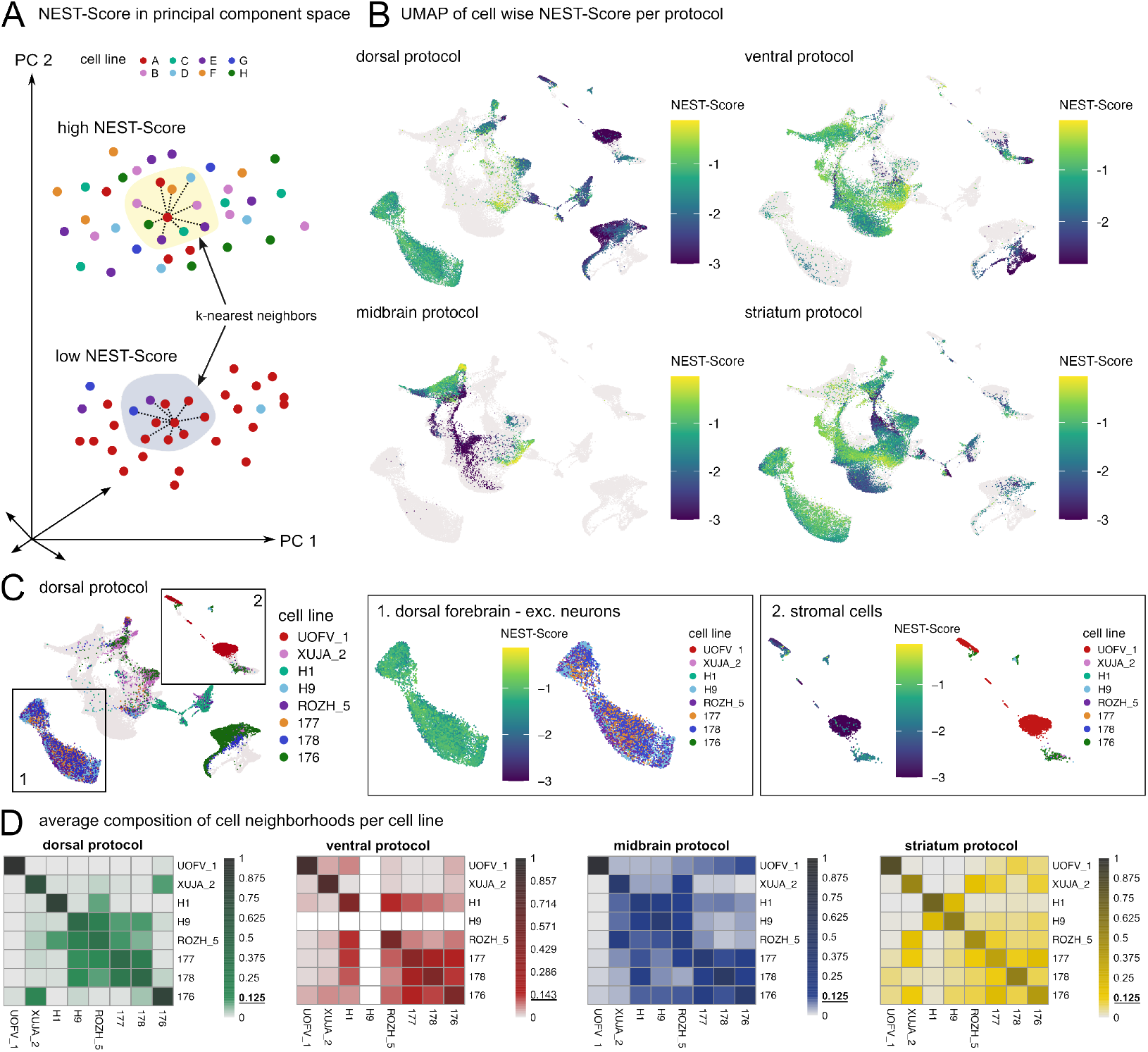
Definition of protocol-driven transcriptional signatures using NEST-scores and cell line biases. (A) Schematic representation of the NEST-Score. High NEST-Scores (close to “0”) per cell are the result of many cells of different cell line origin in a probed cell’s neighborhood (top panel), while low (negative) NEST-Scores are the result of a probed cell’s neighborhood consisting predominantly of cells of the same cell line origin. (B) NEST-Score distribution within one protocol depicts protocol-driven cell groups that were consistently developed across multiple cell lines. (C) Zoom-in dorsal protocol-driven excitatory differentiation lineage (1.) and not-well-mixed muscle cells (2.) in dorsal protocol. (D) Scaled cell line frequencies (a cell’s local neighborhood cell line distribution divided by the global cell line frequency) average across cells of a cell line show that cell lines mix well with different cell lines for each specific protocol. In a perfect mix cell lines would contribute to 1*/*8 = 0.125 of a cell’s scaled neighborhood for eight cell lines (dorsal, midbrain and striatum protocol) and 1*/*7 ≈ 0.143 for the seven cell lines in the ventral protocol.

In more detail, for each protocol, we performed Principal Component Analysis (PCA) on all cells originated in that protocol. Then, for each cell, we determined the cell line origins of its 100 nearest neighbor cells (Fig. S2A, Methods) considering the top principal components also used for downstream analysis (see Methods). This results in a cell-wise cell line frequency vector which then was compared to the global cell line frequency vector utilizing the negative KullbackLeibler divergence (31) (Methods). The NEST-Score reaches its upper bound, “0”, when both frequency vectors agree and hence the respective cell has a neighborhood consisting of all considered cell lines (Fig. 2A, Fig. S2A, Methods). By applying a threshold (design dependent on number of cell lines, see Methods) to the NEST-Scores we can classify each cell in downstream analyses as either protocol-driven (high NEST-Score) or cell line-driven (low NEST-Score). Plotting the NEST-Scores of each cell onto the protocol-resolved UMAPs (Fig. 2B, C) allowed us to visualize how reliably different cell lines generate protocol-driven cell types within each protocol. For example, the entire trajectory of dorsal forebrain progenitors, intermediate progenitors and excitatory neurons consists of cells with high NEST-Scores, indicating these cell types are consistently produced across multiple cell lines. Similarly, other clusters on the UMAP are consistently produced across cell lines in other protocols (high NEST-Scores), i.e., interneurons in the ventral protocol. Notably, all protocols also contained some cells in clusters with low NEST-Scores (Fig. 2C), indicating that in such instances the given cell line did not adhere to a given protocol’s guidance cues. These effects vary across protocols and cell lines and cannot be explained by assuming that guidance cues of a protocol affect cell lines uniformly. For example, cell line 176 produced mostly protocol-driven cells in ventral, midbrain and striatum protocols, but not in the dorsal protocol, whereas Uofv_1 predominantly produced cell line-driven cell types across all four protocols (Fig. S1E, Fig. S2B, Fig. S2C). This analysis enabled us to evaluate the performance of all cell lines in all protocols individually and to compare them pairwise to identify cell lines that act similarly. For example, we observed that the four cell lines H9, Rozh_5, 177, and 178 produced similar cell states throughout the dorsal protocol (Fig. 2D), indicating that cells derived from these cell lines were commonly guided by protocol cues. In contrast, cell lines such as H1, Uofv_1, Xuja_2 and 176, predominantly mixed only with themselves, consistent with these lines being cell line-driven in this particular protocol. Analyzing all cell lines in all protocols, we find that individual cell lines may have cell line-driven biases for cell generation in different protocols, but most growth conditions support the generation of desired protocol-driven cells across cell lines.

The use of the NEST-Score discussed above is highly depen-dent on which cell lines are considered for analysis. When multiple cell lines exhibit similar cell-line driven biases, cells – and subsequently cell clusters – may be classified as protocol-driven. Therefore, we asked whether protocol-driven cell states and clusters are also the ones that occur in protocol’s target tissue as annotated by region and cell type (compare Fig. 1B) (13, 20, 27, 29). When aggregating the NEST-Scores by annotated cell type, we found that all main glial and neuronal cell type clusters visualized in the pan-protocol, pan-cell line UMAP have high scores for the cell types intended by the respective protocol (Fig. 2B, Fig. S2C). For example, In the dorsal protocol, dorsal progenitors and excitatory neurons were generated across multiple cell lines. In the ventral protocol, instead, ventral progenitors and inhibitory neurons are protocol-driven, and in the midbrain protocol midbrain progenitors and neurons are protocol-driven. In contrast, muscle and stromal cells that are found in the dorsal protocol have a low NEST-Score, indicating they were mostly generated by single cell lines (in the dorsal protocol 79.3% of stromal cells derived from cell line Uofv_1 and 86.40% of muscle cells from cell line 176) (Fig. 2C, Fig. S2B, Fig. S2C). Our analysis also identified cell states that are protocol-driven in one protocol, but cell line-driven in another protocol underlining the necessity for comparing four different protocols. For example, while most cell lines produced protocol-driven cells in the midbrain protocol, one cell line, Uofv_1, generated interneurons in this protocol (Fig. 2B, Fig. S2B). Cells of the interneuron lineage on the other hand are protocol-driven in the ventral and striatum protocols.

By defining protocol-driven cell states generated from multiple cell lines across four brain organoid protocols and providing the NEST-Score as a metric to score the reliability of such cell generation we provide a new protocol benchmarking analysis scheme. The definition of protocol-driven cells per growth protocol thus comprises the first success factor in the application of a robust brain organoid protocol. Importantly, this method can be used to evaluate the suitability of any additional cell line for growing brain organoids by recomputing the NEST-Score based on scRNA-seq data obtained using this cell line.

### Comparison of brain organoid potential to fetal references

Our data allows us to distinguish cell line-driven cell types unique to one cell line from cell types which are protocol-driven and consistently generated in a given protocol. To test, how well the protocol-driven cell types in our four protocols cover the diversity of cells found in the human brain, we compared them to two commonly referenced *in vivo* fetal brain tissues atlases (23, 24). We used a dataset combining individual forebrain, midbrain and hindbrain as well as separate dorsal and ventral preparations (24) and a second dataset containing cortical regions that has previously been compared to brain organoids (23). Since both *in vivo* dataset comparisons gave comparable results (Braun et al. (24) comparison Fig. 3A-F and Fig. S3A, B; Bhaduri et al. (23) comparison Fig. S3C-H), we focused on the dataset including more brain regions (24).

**Fig. 3.**
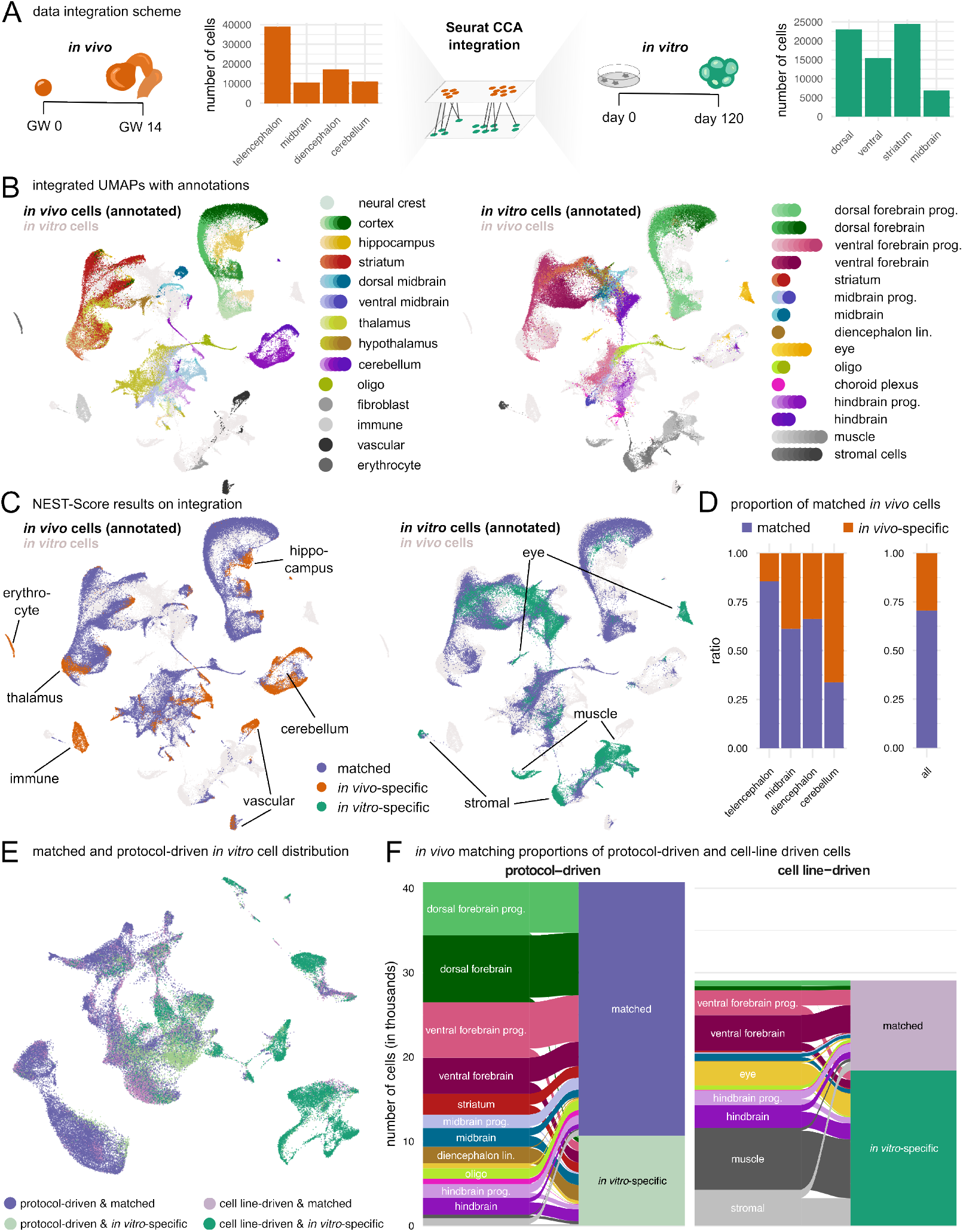
Data integration of *in vitro* derived orgnoid cells with *in vivo* brain references. (A) Cell numbers of the *in vivo* dataset split by sequenced brain region and cell numbers of the *in vitro* dataset split by protocol. Schematic of Seurat CCA integration of *in vivo* fetal brain samples from gestational week (GW) 14 (24). (B) Sequenced regions of *in vivo* and annotated, region-specific cell types of *in vitro* datasets align in the integrated UMAP. (C) *In vitro* / *in vivo* overlapping and non-overlapping regions colour-coded on the integrated UMAP. Cell types with low *in vivo* / *in vitro* overlap are indicated. (D) Proportions of *in vivo* / *in vitro* overlapping cells of all *in vitro* cells split by *in vivo* annotations. (E) Integrated *in vivo* / *in vitro* UMAP coloured by groups of protocolor cell-line driven *in vitro* cells for their overlap with *in vivo* reference. (F) Proportions of protocol- and cell line-driven cells split by cell types falling into *in vivo*-mixed and not *in vivo*-mixed categories.

We first subset the fetal reference to gestational week (GW) 14, the time point most closely resembling day 120 brain organoids (32). Integrating the *in vivo* reference with our pan-cell line, pan-protocol dataset using Seurat’s CCA algorithm (33) (Methods, Fig. 3A) revealed a large overlap as visualized in the resulting UMAP (Fig. 3B, Sup. Table 2). Clusters annotated as dorsal forebrain in organoids *in vitro*, for example, overlap with cortex clusters *in vivo*. Remaining residual discrepancies between the *in vivo* and *in vitro* data are expected and indicate that biological variation is not over-corrected.

To quantify the overlap between *in vivo* and *in vitro* in a systematic manner, we again applied the NEST-Score workflow to access the contribution of *in vivo* and *in vitro* samples to a cell’s neighborhood (Fig. 3C, Methods). We found that substantial portions of cell types in the integrated dataset, were covered by both *in vitro* organoid and *in vivo* fetal cells (Fig. 3D). Thereby, such cells fulfill the second success criteria for brain organoid growth in that they resemble *in vivo* counterparts. Importantly, while each protocol individually allows only limited coverage of the fetal tissue diversity (dorsal protocol covers 41.4%, ventral 50.8%, midbrain 27.3% and striatum 61.2% of the cells in the *in vivo* reference data set) (Fig. S3A), combining the four *in vitro* protocols resulted in an overall 70.6% representation of all fetal brain cell states (Fig. 3D). The remaining 29.4% of the *in vivo* cells that do not have *in vitro* cells in their neighborhood include immune cells, erythrocytes and vascular cells (Fig. 3B, C), cell types originating outside the brain that are known not to be included in brain organoids. We also did not cover all cerebellar cell states, possibly because hindbrain- or cerebellum-specific brain organoid protocols were not included in our study.

We sought to combine our two success factors for organoid growth, (1) generation of protocol-driven cells across cell lines in each protocol and (2) the generation of *in vivo* matched cell types. In this analysis we observed that most protocol-driven *in vitro* cells (namely the major dorsal, ventral and midbrain glia and neuron populations) also have counterparts *in vivo* (Fig. 3E, F). This indicates that successful adherence to protocol cues results predominantly in cell states present *in vivo* being produced, highlighting the robustness of protocol characterization across multiple cell lines for evaluation of *in vivo* cell type generation. In contrast, most cell line-driven cells derived from cell lines not adhering to protocol cues, do not mix with any cells from the *in vivo* references. As expected, these include muscle and stromal cells (Fig. 3E, F, Fig. S2C, Fig. S3B).

### Bulk RNA-Sequencing Derived, Time-Resolved Transcriptional Signatures

Our scRNA-seq analysis was confined to day 120, a time point where a large variety of both progenitor and differentiated cell types can be observed. To test, whether organoid development at earlier time points can predict later cell stages, we performed a time-course of bulk-RNA sequencing experiments to find predictive markers. For all combinations of the four protocols and seven cell lines we sequenced three replicate organoids at days 13/16, 25, 40, 80 and 120 (Fig. 4A, Fig. S4A). We also sequenced three samples of each pluripotent stem cell line in the pluripotent state (day 0). PCA analysis of the combined datasets showed a gradient of samples according to sampling time in PC1 and PC2 (Fig. 4B), while PC3 and PC4 show grouping per protocol (Fig. 4C). Interestingly, grouping per cell line was only visible in PC7 and PC8 (Fig. S4B). By quantifying the contribution of each experimental condition to the total variation of the data (34, 35) (Methods), we found that the sampling time point explains 39.04% of the overall variance while protocol and cell line choice contribute 13.29% and 8.05% to the overall variance observed, respectively (Fig. 4D). Protocol and cell line induced variability was consistent over time (Fig. 4E). In the pluripotent state (day 0), in the absence of protocol cues, cell line identity accounts for more than half of the variation in the data (Fig. 4E). We observed that some cell lines display a more similar expression profiles at this stage (i.e. H1 with Xuja_2 or 177 with 178) (Fig. 4F). However, the overall pairwise correlation between all cell lines is very high (Fig. 4G) and the cell line grouping is not predictive for common organoid outcomes at later stages (Fig. 4H, Fig. S4C-F). This indicates that the transcriptome of pluripotent cell lines cannot be used to predict biases during organoid differentiation. Beginning at day 40 we observed a protocol-specific separation of sample-wise expression profiles (Fig. 4I). At that time point we also found that cell lines mainly producing protocol-driven cells largely cluster together (compare Fig. S2B and Fig. S4C-F). However, this prediction does not hold 100% true for all cell lines in all protocols, i.e., in one case for the dorsal protocol, cell line similarities are different at day 40 compared to 120, showing H9 as an outlier cell line at day 40, and 176 as outlier cell line at day 120 (Fig. 4I). The latter meets the observation of scRNA-seq samples at day 120, where H9 produces abundant protocol-driven cells while 176 does not.

**Fig. 4.**
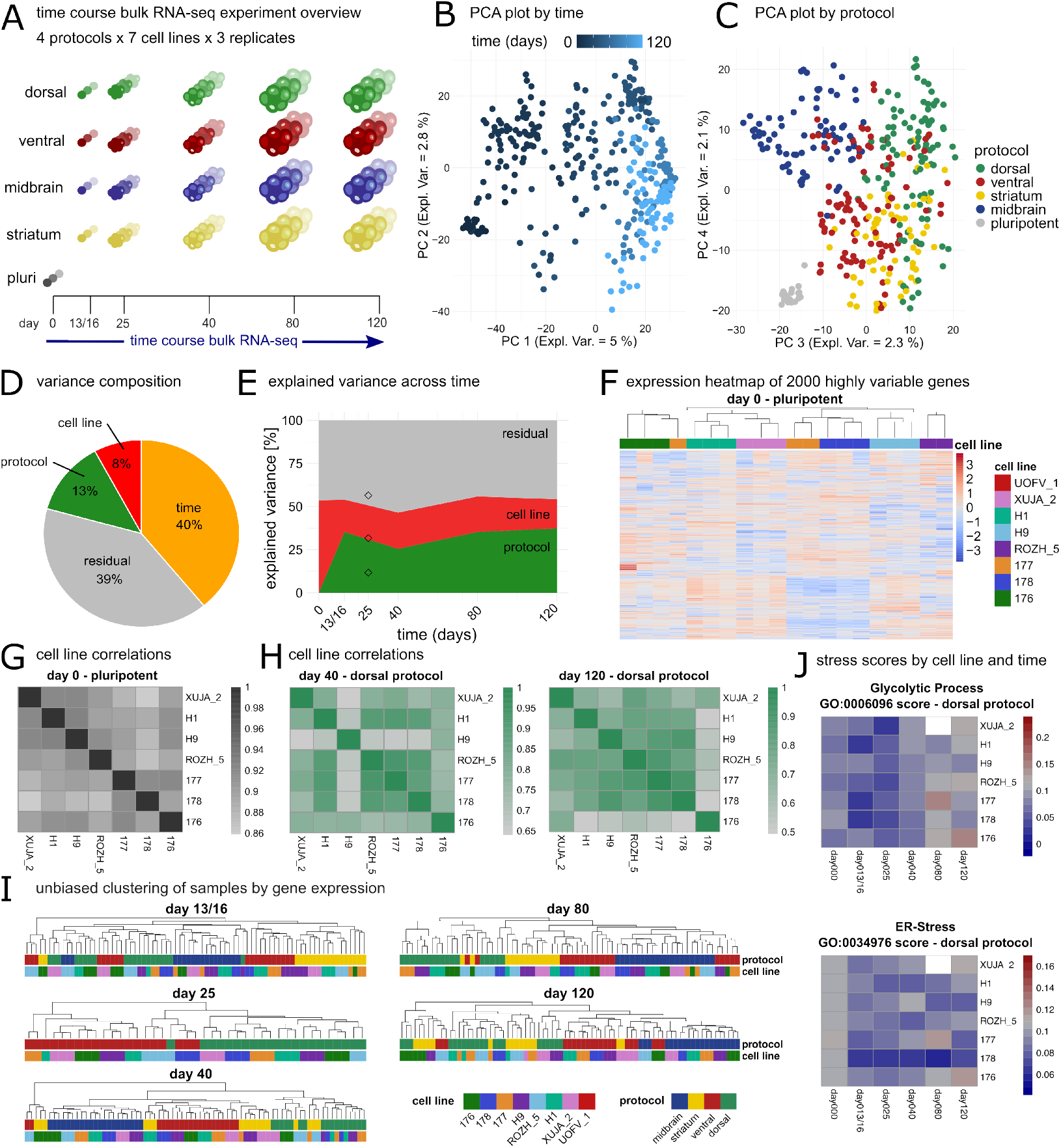
Time-course bulk RNA-seq analysis of brain organoids reveals differences of variation across time, protocol and cell line. (A) Experimental overview for time-course bulk RNA-seq experiment, abbreviation pluri. = pluripotent. (B) Principal component analysis (PCA) of bulk RNA-seq datasets. Plotted are principal components 1 and 2 with samples colored by organoid age. (C Principal component analysis (PCA) of bulk RNA-seq datasets. Plotted are principal components 3 and 4 with samples colored by growth protocol. (D) Overall experimental variance split by factors time, protocol, cell line and residual. (E) Explained variance split by protocol, cell line and residual across time. (F) Hierarchically clustered expression heatmap of 2000 most highly variable genes on day 0 (pluripotent stem cell stage). (G) Pearson correlations between cell lines at day 0. (H) Pearson correlations between cell lines at day 40 and 120 in the dorsal protocol. (I) Hierarchically clustered expression grouping based on 2000 most highly variable genes on days 13/16, 25, 40, 80, 120. (J) Stress related Gene Ontology gene group expression over time in cell lines.

We wanted to understand whether there are communalities between cell lines whose intrinsic biases prevent the production of protocol-driven cells. Given that different cell line-driven cell lines produced different cell line-driven off-target cells, we asked if there is nonetheless a common feature among cell lines unable to produce protocol-driven cell types at day 120. Many organoid studies describe a connection between increased cellular stress levels and decreased similarity of organoid cells to *in vivo* data (20, 36, 37). Organoid stress may result from growth media conditions and nutrient limitations due to missing vascularization (38, 39). We asked if increased stress less levels might be a correlative with cell line-driven tissue growth. To this end, we analyzed the bulk RNA-seq samples over time with respect to high expression of genes assigned to Gene Ontology (GO) terms ‘Glycolytic Process’ (GO:0006096) and ‘response to endoplasmic reticulum stress’ (GO:0034976, ER-stress) indicative of a stress response (Methods). ER-stress is readily apparent in the pluripotent stem cell stage at day 0 and decreases after the application of a growth protocol in the first 40 days of organoid growth, consistent with previous data (40). At later time points, both stress scores increase again, but to different extents across cell lines. Indeed, at day 120 in the dorsal protocol high stress scores are observed for cell lines 176 and XUJA, which generated mainly cell line-driven tissue (Fig. 4J, Fig. S2A), suggestive of a limiting effect of cellular stress in cells to respond to protocol cues at late stages of organoid development.

### Protocol-specific markers for successful organoid growth

The time-resolved bulk RNA-sequencing data allowed us to identify protocol-specific markers independent from individual cell lines. We sought genes that are highly expressed across all cell lines in one protocol compared to all other protocols and time points (Fig. 5A, Sup. Table 3). Reassuringly, marker genes identified in this way match expression patterns of protocol targeted cell types in day 120 scRNA-seq data (Fig. S5A). However, this list also includes markers for earlier time points, like SIX6 for the ventral protocol (Fig. 5A, B), GLI3 for the dorsal protocol (41) and EN1 and FOXA2 for the midbrain protocol (42). Besides transcription factors (highlighted in Fig. S5B) we also found developmental morphogens (i.e. WNT2B for dorsal protocols) or morphogen receptors (i.e. PTCH1 in the ventral protocol) and ion channels (i.e. KCNL13 and TRPM3 in the dorsal protocol) (Fig. 5A). In a second analysis, we concentrated our search for genes that are highly expressed at day 40 in cell lines that produce protocol-driven high-quality organoids at day 120 and compared them to the other protocols. This results in a marker gene list that is consistent with successful organoid derivation and may serve as a simple quality control readout via RNA-sequencing or quantitative real-time PCR of individual organoid batches months before organoids have fully matured. Day 40 was chosen since this is the earliest time point in which we observe consistent sample separation according to protocol. For each protocol, we tested cell lines yielding a minimum of 75% protocol-driven cells (compare to Fig. S2B) against the remaining cell lines in this protocol and all cell lines in the other three protocols (Fig. S5C, Sup. Table 3). The resulting list of upregulated genes again contains expected marker genes (i.e. EOMES, TBR1 in the ventral protocol and DLX genes in the ventral protocol) alongside additional putative protocol specific early regulators of successful organoid derivation. To make our data easily accessible, we compiled a Shiny App (43) data explorer (https://vienna-brain-organoidexplorer.vbc.ac.at) (Fig. 5C). This allows exploration of gene expression signatures for all considered protocols and cell lines across bulk RNA-seq timepoints and comparisons of *in vivo* and *in vitro* scRNA-seq datasets. Furthermore, it can be used to analyze expression patterns of entire gene sets based on their GO term association, suggesting, which biological processes may be important in specific cell types at specific time points and in specific protocols.

**Fig. 5.**
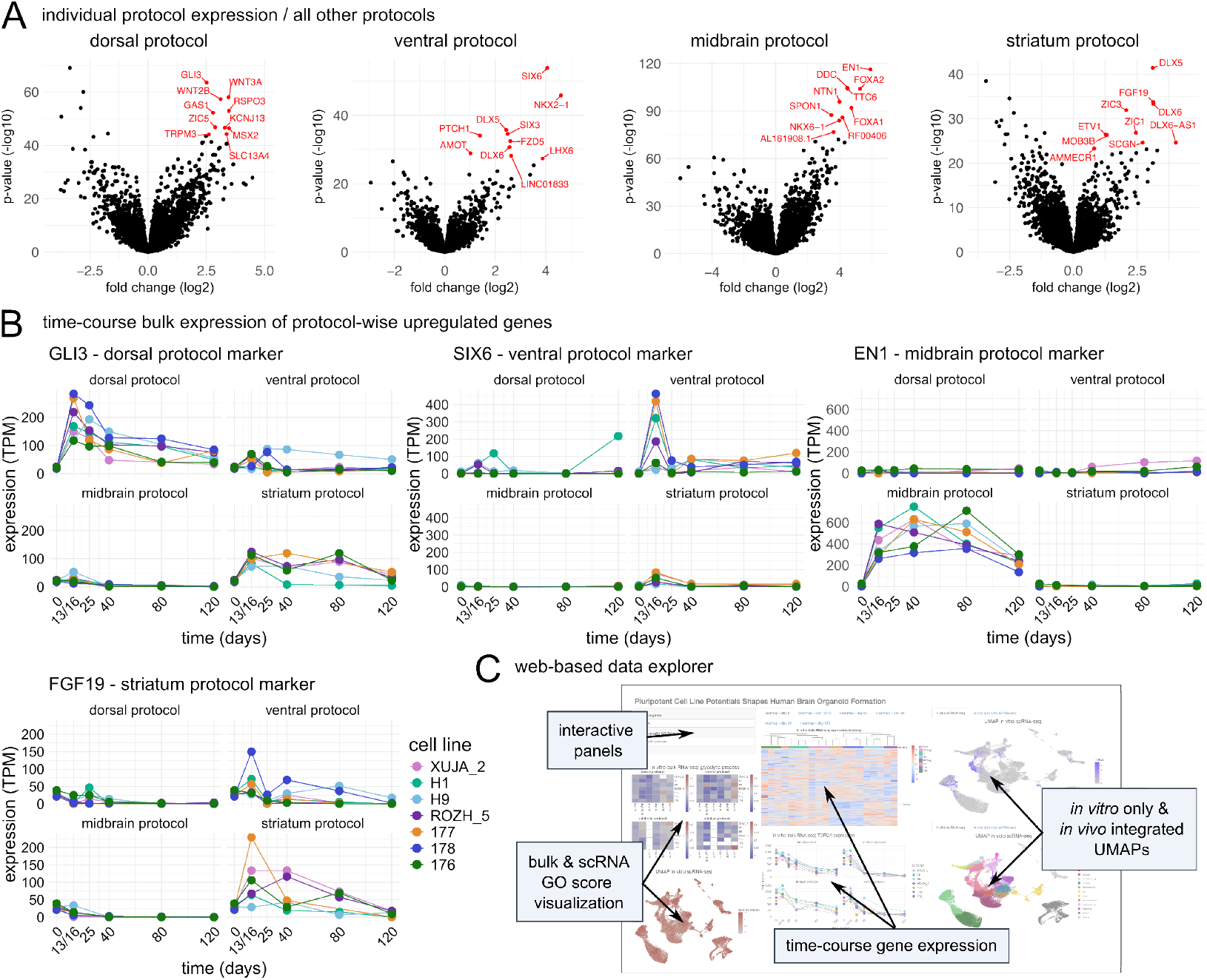
Time-course bulk RNA-seq identifies protocol specific signatures and data explorer overview. (A) Volcano plots of differentially expressed genes tested against all remaining three protocols separated by indicated protocol. Top 10 upregulated genes are indicated. (B) Gene expression in transcripts per million (TPM) of exemplary protocol markers shown for each cell line over time. Protocols and marker genes are indicated. (C) Screenshot of interactive data browser that allows users to browse through expression profiles of genes of interest in both time course bulk RNA-seq and UMAPs of scRNA-seq data (*in vitro* only and *in vivo* integrated) as well as user-defined Gene Ontology analysis.

## Discussion

Brain organoids are widely used in neurodevelopment research and disease modeling (1, 2), but variability across cell lines or protocols makes their use challenging. Here, using only four protocols we provide evidence for generation of large portions (70.6%) of fetal brain cells across several pluripotent stem cell lines. Using multiple cells lines allows to define the propensities of brain organoid protocols for cell modeling independent of individual cell line tested, providing a valuable reference for future organoid derivation from additional cell lines. We provide comprehensive scRNA-seq and bulk RNA-seq data sets to characterize the four protocols over time and for cell types generated. We introduced the NEST-Score to provide a quantitative measure for reliable generation of cell states across protocols and to evaluate their recapitulation of *in vivo* cell types. Thereby, we define successful protocols by two factors: The generation of (1) protocol-driven cells across multiple cells, that (2) match *in vivo* reference cells. Furthermore, we identified marker gene sets that will allow predictions of organoid protocol success in future work and allow browsing of all our data using an intuitive web portal (https://vienna-brain-organoidexplorer.vbc.ac.at).

By aligning additional cell lines to our reference and *in vivo* data, the NEST-Score can be used to identify protocol-driven cell states for each of the four protocols and to test how well they match their *in vivo* counterparts, respectively. The NEST-Score is particularly useful for the analysis of scRNAseq data that will not require additional batch correction, as we show in the analysis of protocol-driven cell states. When evaluating samples with a considerable batch effect, the NEST-Score may also be applied to batch corrected PCA space, however, in such cases, a reasonable interpretation of the NEST-Score evaluation is only possible, if the strength of the batch effect removal does not remove all, i.e., also biological, variation between samples. Since the NEST-Score defines cell neighborhoods in PCA space, it is independent from non-linear and often information-compromising dimensionality reduction methods such as UMAP. It is calculated cell-by-cell and does not rely on any clustering. Therefore, it is independent of cluster resolutions that might average out cell bias effects. Unlike previous methods (13, 44), it also does not require any prior cell type annotation making it possible to distinguish the success of a protocol based on (1) reliably producing the same cell states and (2) producing the desired cell types. Our approach is motivated by similar strategies, used for example to measure batch effects (45) or to determine a coverage coefficient (20). Since we utilize the Kullback-Leibler Divergence, the NEST-Score has the additional advantage to allow for a combined comparison across more than two conditions (here: multiple cell lines).

Our data show that organoid variability is not just a simple product of protocol stringency and cell line bias. Instead, individual cell lines display distinct biases in a protocol and a protocol’s outcomes should ideally be evaluated from data encompassing a collection of different cell lines, thereby rendering a comprehensive protocol characterization cell line independent. This work is a step towards the goal of recapitulating all brain cells *in vitro*, with a focus on dorsal and ventral forebrain, midbrain and striatum. Given that other brain areas, particularly cerebellum, are not included in our current protocol set, we anticipate that brain cell coverage can even be further increased in the future by adding protocols addressing, e.g., cerebellum, blood vessels and immune cells (46, 47). Our time-course RNA-seq data facilitates the identification of protocol-specific markers that can be tested at early time points in order to predict successful organoid formation, saving time and costs. Our datasets and putative marker lists can also serve as a resource for further optimization of experimental protocols and evaluation of pluripotent cell lines for future research. This may be of particular importance for patient-derived cell lines, which require a solid definition of protocol-driven cells as controls. It also allows us to predict the potential of new cell lines to cover the full spectrum of cell types found *in vivo*. There are several limitations of our study. For example, our current data covers only eight cell lines per protocol. Therefore, future efforts should be geared towards extending the number of cell lines across multiple protocols further. Another limitation is that only four protocols are characterized in-depth. Other studies (14, 22) consider larger number of growth conditions, but use fewer cell lines and earlier analysis timepoints and it will be interesting to cross-compare cell generation across these studies.

Our data using four brain organoid growth protocols across multiple cell lines to generate large portions of cells of the developing human brain may serve as a reference for various applications. To facilitate easy and widespread access to our data we implemented a data explorer that visualizes the expression of protocol markers or GO term specific gene sets over time, protocol and cell lines and also in the day 120 scRNA-seq data. The explorer is available at https://viennabrain-organoid-explorer.vbc.ac.at and allows users to identify marker genes for the formation of specific cell types and as well as to choose the right protocol for a specific scientific question.

## ACKNOWLEDGEMENTS

We thank all members of the Bock, von Haeseler, Esk and Knoblich labs for technical expertise and feedback. We thank IMP/IMBA Biooptics services for expert microscopy and flow cytometry services. We thank M. Gollowitzer and the IMP/IMBA Information Technology Department for IT support. We thank A. Sommer and the VBCF NGS unit (www.viennabiocenter.org/facilities) for consultation and sequencing.

## AUTHOR CONTRIBUTIONS

J.N., M.B., A.v.H., C.E. and J.A.K. designed the study, analyzed data and wrote the manuscript with input from all authors. M.B. performed experiments with help from T.K., S.L. and C.E.. J.N., L.H., M.N., and L.D. performed bioinformatic analysis. Supervision was provided by C.B., A.v.H., C.E. and J.A.K.. C.B., A.v.H., C.E., and J.A.K. acquired funding.

## DECLARATION OF GENERATIVE AI AND AI-ASSISTED TECHNOLOGIES IN THE WRITING PROCESS

During the preparation of this work some authors used ChatGPT (https://chatgpt.com) in order to improve readability of initial text drafts. After using this tool, the authors reviewed and edited the content as needed and takes full responsibility for the content of the published article.

## DATA AVAILABILITY

Requests for further information and resources should be directed to and will be fulfilled by Jürgen Knoblich. This study did not generate new unique reagents. Raw single-cell and time-course bulk RNA-seq data generated and analyzed in this work have been deposited at the European Genome-phenome Archive (EGA), which is hosted by the EBI and the CRG, under series numbers EGAS50000000662 and EGAS50000000663, respectively, and are available through controlled access as of the date of publication. Processed single-cell and time-course bulk RNA-seq data have been deposited on NCBI Gene Expression Omnibus (GEO) via series numbers GSE277968 and GSE277967, respectively, and are publicly available as of the date of publication. R scripts and corresponding source data to reproduce all figures and tables presented in the manuscript are deposited on GitHub (https://github.com/jn-goe/brain_organoids_four_protocols) and Zenodo (DOI: 10.5281/zenodo.13742635) and are publicly available as of the date of publication. Functionalities of the NEST-Score are publicly available as R package on GitHub (https://github.com/jn-goe/NESTScore) and on Zenodo (DOI: 10.5281/zenodo.13974435) as of the date of publication. The interactive Shiny App for data exploration is publicly available on https://vienna-brain-organoid-explorer.vbc.ac.at.

## SUPPLEMENTARY TABLES

**Sup. Table 1** Differentially expressed genes across scRNA-seq clustering

**Sup. Table 2** Hierarchical clustering information for *in vitro* and *in vivo* scRNA-seq

**Sup. Table 3** Differentially expressed genes across bulk RNA-seq protocols

## Methods

### Cell Culture

Human embryonic stem cell (hESCs) lines WA01 (H1) and WA09 (H9) were obtained from WiCell (https://www.wicell.org). Human induced pluripotent stem cell lines SCCF-176J clone#1 (abbreviated 176), SCCF-177J clone#8 (abbreviated 177) and SCCF-178J clone#5 (abbreviated 178) were obtained from IMBA iPSC biobank (https://shop.vbc.ac.at/ipsc_biobank). Human induced pluripotent stem cell lines Rozh_5, Uofv_1 and Xuja_2 were obtained from the HipSci consortium (https://www.hipsci.org). All lines were contamination free, STR verified and regularly tested for mycoplasma. Cell lines were maintained according to HipSci recommendations on Vitronectin (Stem Cell Technologies, cat. no. 100-0763) coated plates with Essential 8 Medium (Thermo Fisher Scientific, Catalog # A1517001 or in house produced). All cells were maintained in a 5% CO2 incubator at 37 °C. Cells were either split using DPBS -/-(Gibco, Catalog # 14190-250) or Accutase (Sigma, cat. A6964) and plated in Essential 8 Medium supplemented with RevitaCell Supplement (Thermo Fisher Scientific, Catalog # A2644501).

### Brain organoid generation

Brain organoids were generated as previously described (19, 28). Media compositions are given below. Pluripotent cells were grown to 60-80% confluency and single cell suspensions were obtained using Accutase. Pelleted cells were resuspended in E8 media supplemented with RevitaCell and counted. 8 000-10 000 cells were seeded to form embryoid bodies in a 96-well ultra-low-attachment U-bottom plate (Sigma, cat. #CLS7007).

### Dorsal protoco

Day 0: Seeding in 150 µl E8 supplemented with RevitaCell. Day 3: E8 media. Days 6, 7, 8, 9: Neural induction media (NI), Day 10: 1% Matrigel in NI media (Corning, cat. # 356235) and transfer to 10cm plates coated with anti-adherence rinsing solution (Stemcell Technologies, cat. no. 07010). Day 13, 14: NI media supplemented with 3 mM CHIR (Merck, cat. #: 361571), Days 16, 19, 22: Improved-A media with transfer to shaker on day 19. Days 25-40 every 3-4 days: Improved+A media. Days 40-60 every 3-4 days: Improved+A media supplemented with 1% Matrigel, Day 62: 75% Improved+A media mixed with 25% Brainphys media supplemented with 1% Matrigel. Day 65: 50% Improved+A media mixed with 50% Brainphys media supplemented with 1% Matrigel. Day 69: 25% Improved+A media mixed with 75% Brainphys media supplemented with 1% Matrigel. Days 72-120 every 3-4 days: Brainphys media supplemented with 1% Matrigel, 20 ng ml-1 BDNF (Stemcell Technologies, cat. no. 78005.3), 20 ng ml-1 GDNF (Stemcell Technologies, cat. no. 78057.3), 1 mM db-cAMP (Santa Cruz Biotechnology, cat. no. sc-201567C).

### Ventral protocol

Day 0: Seeding in 150 µl E8 supplemented with RevitaCell. Day 3: E8 media. Days 5, 7, 9, 10: Neural induction media (NI) supplemented with 100 nM SAG (Merck, cat. #: US1566660) and 2.5 µM IWP-2 (Sigma-Aldrich, cat. #: 10536) with 1% Matrigel added at day 10. Days 13, 15, 17: Improved-A supplemented with 100 nM SAG and 2.5 µM IWP-2. Organoid maturation from day 19 on as in dorsal protocol.

### Midbrain protocol

Days 0, 2:150 µl NI supplemented with RevitaCell, 200 ng ml-1 Noggin (R&D Systems, cat. no. 6057), 10 µM SB431542 (Stemgent, cat. no. 04-0010-10) and 0.8 µM CHIR. Day 4: NI supplemented with 200 ng ml-1 Noggin, 10 µM SB431542, 0.8 µM CHIR, 300 nM SAG and 100 ng ml-1 FGF-8 (fibroblast growth factor 8; R&D Systems, cat. no. 5057-FF). Day 6: NI supplemented with 300 nM SAG and 100 ng ml-1 FGF-8. Day 8: Improved-A supplemented with 300 nM SAG, 100 ng ml-1 FGF-8, 2% Matrigel. Transfer to a 10cm dish. Day 10: Improved-A supplemented with 2% Matrigel. Day 13: Improved-A. Days 16-25 every 3-4 days: Improved+A media. Organoid maturation from day 25 on as in dorsal protocol.

### Striatum protocol

Days 0, 2:150 µl NI supplemented with RevitaCell, 10 nM SAG, 2.5 µM IWP-2. Day 4: NI supplemented with 10 nM SAG, 2.5 µM IWP-2. Day 6: NI, Day 8: Day 8: Improved-A supplemented with 2% Matrigel. Transfer to a 10cm dish. Day 10: Improved-A supplemented with 2% Matrigel. Day 13: Improved-A. Days 16-25 every 3-4 days: Improved+A media. Organoid maturation from day 25 on as in dorsal protocol.

### Organoid media

#### Neural induction medium (NI)

DMEM/F12 (Invitrogen, cat. no. 11330-057), 1% N2 Supplement (Thermo Fisher, cat. no. 17502001), 1% GlutaMAX-I (Thermo Fisher, cat. no. 35050-038), 1% MEM-NEAA (Sigma-Aldrich, M7145), 1:1000 heparin solution (Sigma-Aldrich, cat. no. H3149-100KU), 1% PenStrep (Sigma-Aldrich, cat. no. P4333).

#### Improved-A medium

50:50 DMEM/F12 : Neurobasal (Gibco, cat. no. 21103049), 0.5% N2 supplement, 2% B27-A (Thermo Fisher, cat. no. 12587010), 1:4000 insulin (Sigma-Aldrich, I9278), 1% GlutaMAX, 0.5% MEM-NEAA, 1% Antibiotic-Antimycotic (Thermo Fisher, cat. no. 15240062).

#### Improved+A medium

50:50 DMEM/F12 : Neurobasal. 0.5% N2 Supplement, 2% B27+A (Thermo Fisher, cat. no. 17504044), 1:4000 insulin, 1% GlutaMAX, 0.5% MEM-NEAA, 1% Antibiotic-Antimycotic, 1% vitamin C solution (40 mM stock in DMEM/F12) (Vitamin C: Sigma-Aldrich, cat. no. A4544), 1 g l-1 sodium bicarbonate (Sigma-Aldrich, cat. no. S5761).

#### Brainphys

BrainPhys Neuronal Medium (Stemcell Technologies, cat. no. 05790), 2% B27+A, 1% N2 Supplement, 1% CD Lipid Concentrate (Thermo Fisher Scientific, cat. no.11905031), 1% Antibiotic-Antimycotic, 1:147 20% glucose solution.

#### scRNA-sequencing

Organoids were dissociated at day 120 by incubation in a 9:1 mixture of Accutase (Sigma Aldrich, cat # A6954) and 10x Trypsin (Gibco, cat # 15400) at 37 °C on a Thermo-shaker (800 rpm) for approximately two hours and four units of TurboDNase (Thermo AM2238) were added after 30 min. After enzymatic dissociation, cells were filtered through a 35 µm strainer followed by dilution with 1% BSA in DPBS-/-(Biostatus; DR70250, 0.3 mM), counted and cells labeled (hashed) with TotalSeq™-A antibodies (biolegend) as required to allow demultiplexing of 4 samples either by genetic background or hash information per single cell transcriptomic reaction. For hashing, cells were stained for 30 min on ice, washed twice with 1% BSA in DPBS-/-. After flow cytometry sorting a pool of equal cell numbers for each of the 4 samples was used as input for the Chromium Next GEM Single Cell 3’ Gene Expression (v3.1) following the 10x Genomics user guide (one experiment of H9 cells in the ventral condition and one sample in 177, dorsal condition failed leading to only three cell lines being loaded in these instances). Sequencing of gene expression and hash libraries was performed on an Illumina NovaSeq S4 lane.

#### scRNA-seq sample pooling and demultiplexing

Sequencing data of 10X libraries was processed using Cell Ranger software (V3.0.2, 10X Genomics) using reference genome GRCh38. Cells were demultiplexed by cell line genotype using Soupor-cell (V2.4) (48) and two replicate organoids per cell line and protocol pooled for analysis. In experiments containing hash information, hash information was used for demultiplexing.

#### Pre-Processing and Downstream Analysis

Per sample resulting cell-by-gene, unique molecular identifier (UMI) count matrices were analyzed in R (version 4.4.0) using Seurat (v5.0.1) (49). We observed high cell-wise expression levels of MALAT1 which is known to be a non-informative sequencing artifact in scRNA-seq data, dominating total number of UMIs in some cells and influencing gene expression normalization and subsequent downstream analysis. Hence, we discarded cells, in which more than 10% of all UMIs were assigned to only one gene and subsequently deleted MALAT1 from the count matrices of all *in vitro* datasets. Then, we filtered for high-quality cells based on doublet detection performed with Python Package ‘scrublet’ (v0.2.3, loaded via R Package ‘reticulate’ (v1.38.0)), number of uniquely detected genes (‘nFeature’) between 500 and 5000, number of UMIs (‘nCount’) between 500 and 10 000, percentage of mitochondrial reads between 0.1% and 15% and percentage of ribosomal reads between 0.1% and 50%.

Counts across all replicates, cell lines and protocols were merged, and genes expressed in less than 10 cells were discarded. After cell-wise count log-normalization (‘LogNormalize’), 3000 highly variable genes were identified (‘FindVariableFeatures’) and ribosomal and mitochondrial genes were removed from the list of highly variable genes. Next, we performed cell-wise cell cycle annotation by Seurat’s function ‘CellCycleScoring’ and gene list ‘cc.gene’. Data was scaled (‘ScaleData’) whilst regressing out cell cycle biases (vars.to.regress = c(‘S.Score’, ‘G2M.Score’) and Principal component analysis was performed (‘RunPCA’). The shared nearest neighbor graph (‘FindNeighbors’) and the Louvain clustering algorithm (‘FindClusters’, with resolution 3.5) as well as the UMAP embedding (‘RunUMAP’) were computed on the first 40 Principal Components. 40 Principal Components were chosen due to flattening of an Elbow Plot showing explained variance against number of Principal Components and because well-known clusters and differentiation trajectories became apparent in the subsequent UMAP visualization. Positive differentially expressed genes per cluster were computed via Seurat’s function ‘FindAllMarkers’ (using the default Wilcoxon Rank Sum test and setting ‘only.pos = TRUE’) (Sup. Table 1). Clusters were annotated based on expression of marker genes and hierarchically aggregated to a coarser cell type annotation (Sup. Table 2, includes top 10 positive differentially expressed genes with p_val_adj < 0.01 and avg_log2FC > 1 for each cluster).

#### VoxHunt correlation analysis

For comparison of the scRNA-seq organoid data with spatial gene expression data on a mouse brain slice (BrainSpan, (26)), we performed VoxHunt (25) with R Package ‘VoxHunt’ (v1.1.0) and used the provided sample of embryonic day 13 (‘E13’). For the Spearman correlation analysis with subsets of the scRNA-seq organoid data, we pooled the top 15 (based on the provided Area under the Curve value, AUC) genes based on the sample’s respective annotation level ‘custom_2’ using VoxHunt’s function ‘structure_markers’.

#### Measure cell neighborhood homogeneity: NEST-Score

The NEST-Score was computed for each protocol separately, hence for one protocol, the data was subset to all cells generated in this specific protocol. The concept of the NEST-Score will be explained with respect to the 8 different cell lines that contribute to the dorsal, midbrain and striatum protocol. For the ventral protocol, only 7 contributing cell lines are present and the different number of cell lines results in a different range of NEST-Scores as can be seen in Fig. 2B.

For each subset, downstream analysis was re-computed with Seurat, in more detail, the computation of the most variable 2000 features (‘FindVariableFeatures’), data scaling (‘ScaleData’), Principal Component Analysis (‘RunPCA’) and nearest neighbor search (‘FindNeighbors’, based on Euclidean distance in the space spanned by the first 40 Principal Components). Subsequently, for a cell *i* and its *k* nearest neighbors 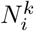, we count how often each cell line *x* occurs. The resulting 8- dimensional vector (since 8 cell lines are considered) is then divided by *k* to obtain the local cell line distribution

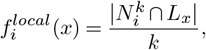

for cell *i* and cell line *x*, where *L*_*x*_ is the set of all cells from cell line *x*. The local cell line distribution vector 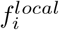 is then entry-wise divided by the global cell line distribution vector *f*^*global*^, i.e., the vector of relative frequencies of cell lines across all cells within the considered protocol:

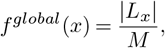

where *M* = Σ*x* |*L*_*x*_| is the number of all cells in the considered protocol. This yields the scaled local cell line distribution

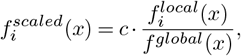

for cell *i* and cell line *x*, where *c* is a constant such that 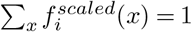.

Finally, the NEST-Score *N* (*i*) for cell *i* is defined as negative Kullback-Leibler divergence (31) of 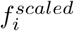 and the 8-dimensional uniform frequency vector *f*^*uniform*^ = (1*/*8, …, 1*/*8):

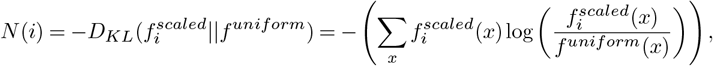

for cell *i*.

The minus provides a more intuitive interpretation: In case the neighborhood of a cell is perfectly resembled by all considered cell lines, the local cell line distribution coincides with the global cell line distribution, yielding 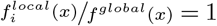 for all cell lines *x*. Accordingly, 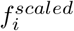 becomes a uniform distribution and hence also 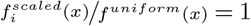 for all cell lines *x*. Since 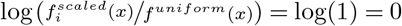, the NEST-Score reaches its upper bound 0 in this scenario. In our application we interpret this as the expression state of (and around) cell i is produced by multiple cell lines and hence consistently recovered in the considered protocol.

To binarily classify a cell as either protocol-driven (= well-mixed neighborhood with respect to cell line abundances) or cell line-driven, we computed a NEST-Score threshold as following: As a minimum requirement to identify a cell state as protocol-driven, we want to observe at least three different cell lines in the respective cell’s neighborhood. Hence, we compute the NEST-Score for an artificial scaled local cell line distribution 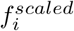, which would appear if a cell’s neighborhood would only consist of two cell lines with equal frequencies, i.e.,

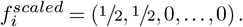

For the eight cell lines (in dorsal, midbrain and striatum protocol), this results in the threshold of 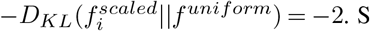. Since only 7 cell lines contribute to the ventral protocol, the threshold computation results in a value of around -1.8. A cell is then classified as protocol-driven, as soon as its NEST-Score is strictly greater than the protocol-specific threshold.

For all presented analysis, we choose *k* = 100 neighbors, since we observe a decreasing NEST-Score for *k* ∈ {1, …, 99} and a more or less constant NEST-Score for *k* ∈ {100,…, 200}. This is shown in Fig. S2A for the dorsal protocol, where we compute the NEST-Scores across multiple *k*∈ {1, …, 200} for each cell, resulting in cell-wise NEST-Score lines. Those lines are colored according to our binary assignment of cells as cell line or protocol-driven based using *k* = 100 and the above introduced NEST-Score threshold of -2.

For the analysis of mixedness between *in vivo* and *in vitro* datasets, we consider the two sample conditions *in vivo* and *in vitro* instead of different cell lines. Again, the computation of a cell’s 100 nearest neighbors was performed in Principal Component Space after batch correction (50 Principal Components to be consistent with the downstream analysis of the integrated data). For the protocol-wise analysis, the integrated data was subset to *in vivo* and *in vitro* data of only the considered protocol. Since during Seurat CCA integration expression matrices are already subset to highly variable features, we only re-computed the Principal Component Analysis and subsequent nearest neighbor search (‘FindNeighbors’, based on Euclidean distance in the space spanned by the first 50 Principal Components) for the respective, batch-corrected subset. For a similar binary classification scheme as above, we computed a NEST-Score threshold based on the scenario that a cell’s scaled neighborhood only originates from one of the two conditions. In more detail we compute the NEST-Score using

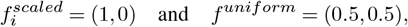

resulting in value of 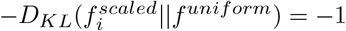, which is also the minimal NEST-Score that can be attained in this setting. If a cell’s NEST-Score was strictly larger than this threshold, i.e., if at least one cell from each condition was present in sthe neighborhood, the cell was labeled as matched. Otherwise, it was labeled as *in vivo* / *in vitro* specific, depending on whether the cell is of *in vivo* or *in vitro* origin.

#### Module Score based on bulk RNA-seq differentially expressed marker genes

Per protocol, the first 50 differentially expressed genes (ranked by adjusted p-value) based on the bulk RNA-seq data across all considered experimental time points were input into Seurat’s ‘AddModuleScore’ function (50) which calculates their average expression and rescales it with respect to a random control gene set.

#### Gene Ontology Scores for scRNA-seq

For one Gene Ontology term, genes attributed to this term were downloaded using the function ‘select’ of R package ‘AnnotationDbi’ (version 1.66.0) and package ‘GO.db’ (version 3.19.1) and input into Seurat’s ‘AddModuleScore’ function.

#### Integration of in vivo and in vitro scRNA-seq data

For fetal brain data published in (24), we merged already quality controlled samples that were sequenced in Gestational Week 14 without any batch correction procedure as provided on https://github.com/linnarsson-lab/developing-human-brain (last access on July 16th 2024). Then, we followed a similar downstream analysis as for the *in vitro* data. We identified the most 3000 highly variable genes, removed ribosomal and mitochondrial genes and MALAT1 from this list. Next, the *in vivo* and *in vitro* data were integrated using Seurat CCA: 1. Integration features (‘SelectIntegrationFeatures’, excluding mitochondrial and ribosomal genes and MALAT1) and subsequently integration anchors (‘FindIntegrationAnchors’, based on canonical correlation analysis ‘CCA’) were determined. After integrating the data with ‘IntegrateData’, downstream analysis was performed as described above to obtain the *in vitro* pan protocol and pan cell line UMAP embedding using the default of all 50 Principal Components. For visualization of clusters, we combined information of metadata ‘Subregion’ and ‘CellClass’ (Sup. Table 2). The alluvial plot in Fig. 2F was plotted with R packages ggplot2 (v3.5.1) and ggalluvial (v0.12.5).

For fetal brain data published in (23), we integrated high-quality cells from Gestational Weeks 14, 17 and 18 and excluded cells not listed in or labeled as ‘Outlier’ in the provided metadata. We randomly (stratified by cell type) subsampled 46 171 cells from the around 205 000 high-quality cells in Gestational Week 18, such that all considered *in vivo* cells in total add up to 70 000 cells roughly matching our *in vitro* data set size. The *in vivo* was integrated with our *in vitro* data as described above, but due to visible batch effects between individuals, we split the considered *in vivo* data per individual (14, 17, 18, 18_2) and treated the respective 4 datasets as separate batches. Integration Anchors (‘FindIntegrationAnchors’) were then computed by setting *in vitro* data as reference dataset. For visualization of clusters, we combined information of metadata ‘structure’ and ‘cell_type’ (Sup. Table 2). For a straight-forward comparison with our *in vitro* dataset, brain region and cell type annotations of both *in vivo* datasets were aggregated hierarchically and renamed into simpler and coarser annotation names (Sup. Table 2).

#### Bulk RNA-sequencing

RNA samples were extracted from organoids collected in Buffer RLT (Qiagen, cat. # 79216) using the RNA isolation kit provided by VBC core facilities. The kit uses carboxylate-modified Sear-Mag Speed beads and was applied using the Kingfisher instrument (Thermo). For RNA sequencing Lexogen’s Quantseq kit was used, including the UMI extension (Lexogen, cat# 015.384, 081.96). Sequencing was performed on Illumina NextSeq High Output 75 cycle lanes and Illumina Novaseq S1 100 cycle lanes and reads combined. All kits were used according to manufacturers’ instructions. Samples were collected at crucial times during protocols (days 13, 25, 40, 80, 120 for dorsal and ventral, and days 16, 40, 80, 120 for midbrain and striatum).

#### Time course bulk RNA-seq analysis

Reads were preprocessed using umi2index (Lexogen) to add the UMI sequence to the read identifier, and trimmed using BBDuk v38.06 (ref=polyA.fa.gz,truseq.fa.gz k=13 ktrim=r useshortkmers=t mink=5 qtrim=r trimq=10 minlength=20). Reads mapping to abundant sequences included in the iGenomes UCSC hg38 reference (human rDNA, human mitochondrial chromosome, phiX174 genome, adapter) were removed using bowtie2 v2.3.4.1 alignment. Remaining reads were analyzed using genome and gene annotation for the GRCh38/hg38 assembly obtained from Homo sapiens Ensembl release 94. Reads were aligned to the genome using star v2.6.0c, alignments were processed using collapse_UMI_bam (Lexogen) and reads in genes were counted with featureCounts (subread v1.6.2) using strand-specific read counting (-s 1).

#### Pre-Processing and Downstream Analysis

For all bulk RNA-seq analysis, R version 4.4.0 was used. To address variable sequencing depths across samples we included only samples with more than 100 000 reads as high-quality samples in the analysis of the bulk RNA-seq data. For PCA read counts were normalized for library size to transcripts per million (TPM) and log2 transformed. Then the 2000 most variable genes were selected and PCA was performed using the ‘prcomp’ command from the R package ‘stats’ (version 4.4.0).

#### Explained variance by experimental condition

To understand how much the individual factors influence gene expression, we first normalized the count data using the ‘voom’ function from limma (version 3.60.0) (35). With the R package ‘variancePartition’ (version 1.34.0) (34) and its function ‘fitExtractVarPartModel’ we then obtained estimates for how much variation of each gene was explained by the individual factors time, protocol and cell line. We multiplied the proportional explained variance of each gene for the different factors with the variance of the respective genes across the samples resulting in the fraction of explained variance per gene. Finally, to gauge the total impact of each factor on gene expression, we added up these individual contributions (fractional explained variance) per factor for all genes. For computing the explained variance of the factors over time we did the same as described above only for each timepoint individually.

#### Gene expression heatmaps and sample clustering

For creating the heatmaps of the expression data counts were variance-stabilized using the ‘vst’ function from DESeq2 (version 1.44.0), then mitochondrial and ribosomal genes were excluded from the variance-stabilized data and the plots created using the ‘pheatmap’ package (version 1.0.12) based on the 2 000 most variable genes.

#### Gene Ontology Scores for bulk RNA-seq

Bulk samples were stored as a merged Seurat Object and genes were retained if expressed in more than 10 samples. Default downstream analysis was performed (treating one bulkRNA-seq sample as one ‘cell’), i.e., log-normalization, determination of 2 000 highly variable genes and scaling. For one Gene Ontology term, genes attributed to this term were downloaded using the function ‘select’ of R package ‘AnnotationDbi’ (version 1.66.0) and package ‘GO.db’ (version 3.19.1) and input into Seurat’s ‘AddModuleScore’ function.

#### Protocol-specific marker genes

In order to find specific genes which are uniquely or predominantly expressed in each experimental protocol, we performed differential expression analysis. Using the R package limma (version 3.60.0) we tested the expression of each gene in a given protocol across all cell lines against the average expression of the gene in all other protocols. We performed moderated t-tests, corrected the p-values via the Benjamini-Hochberg method and only considered genes with a log fold change larger than 0.2.

## Supplementary Figures

**Fig. S1.**
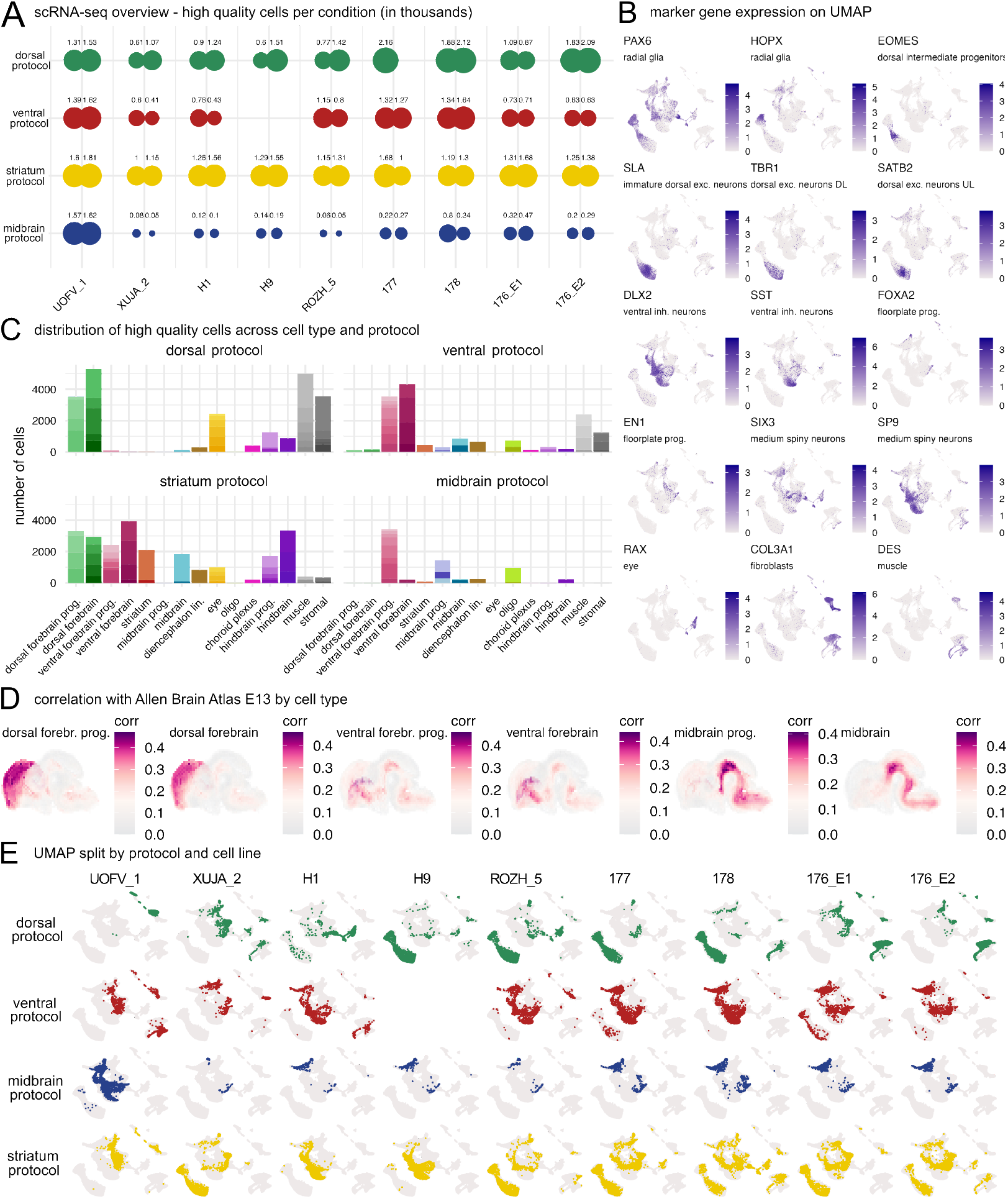
The transcriptional potential of organoids grown from multiple protocols and cell lines. (A) Detailed experimental overview for endpoint scRNA-seq analysis. Analyzed number of organoids and high-quality cells per experimental condition for the scRNA-seq study. (B) Exemplary marker gene expressions on pan protocol pan cell line UMAP. Abbreviations: exc. = excitatory, inh. = inhibitory, prog. = progenitors. (C) Cell type abundances across all sequenced cells and split by annotated cell types. Abbreviations: lin. = lineage, prog. = progenitors. (D) VoxHunt (25) Spearman correlation analysis with respect to annotated cell types of the combined scRNA-seq dataset. (E) Distribution of cells on the combined UMAP split by cell line and protocol as indicated.

**Fig. S2.**
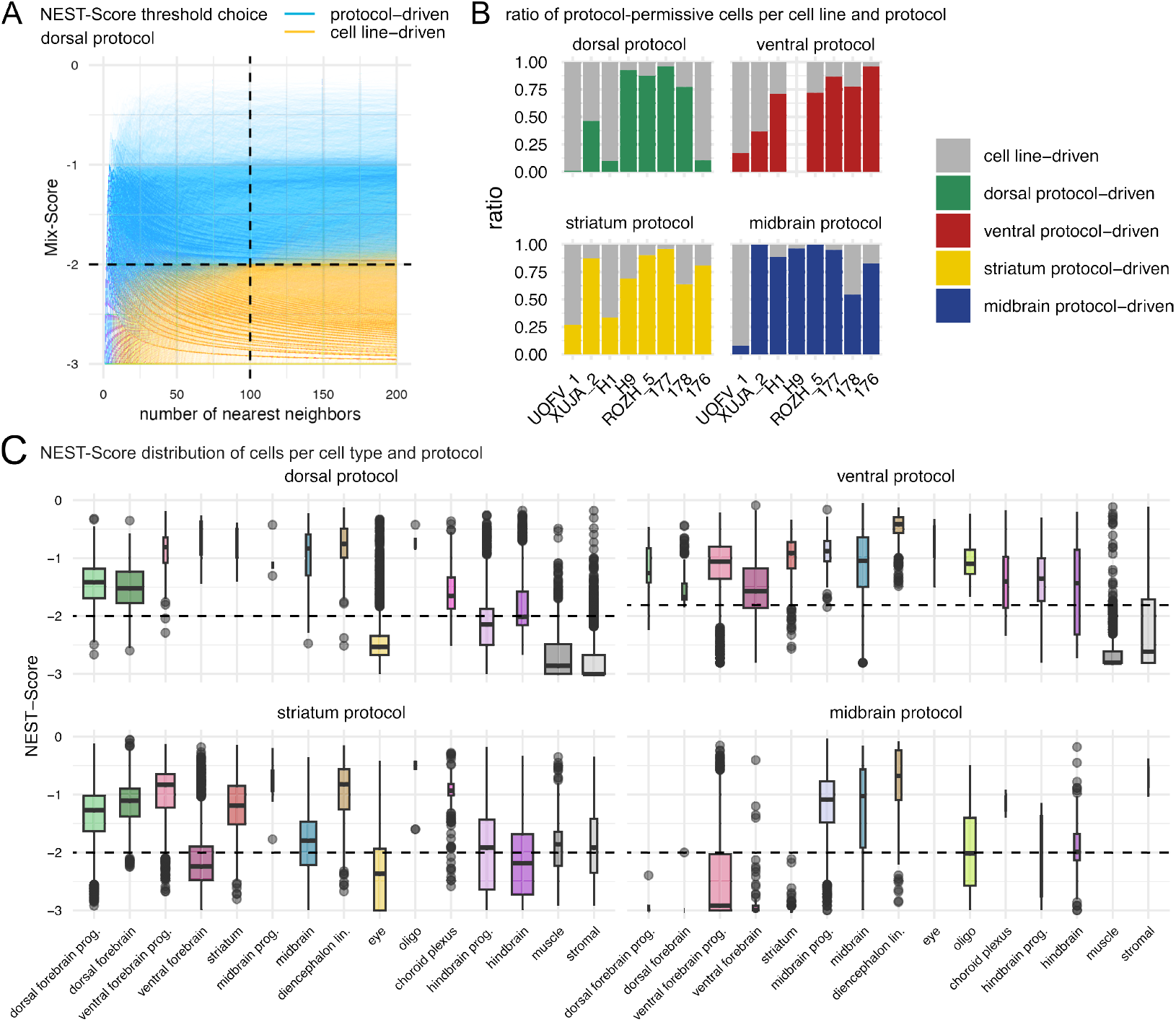
Analysis of protocol-driven cell types and cell line potential. (A) NEST-Scores for dorsal cells computed for a range of nearest neighbors. Protocol- and cell-driven cells are color-coded based on a threshold of -2 (Methods) for 100 nearest neighbors. (B) Number of protocol-driven cells per cell line and protocol as indicated. (C) NEST-Score distribution across cell types and protocols. The width of boxplot scales with the number of cells in the respective cell type, abbreviations: lin. = lineage, prog. = progenitors.

**Fig. S3.**
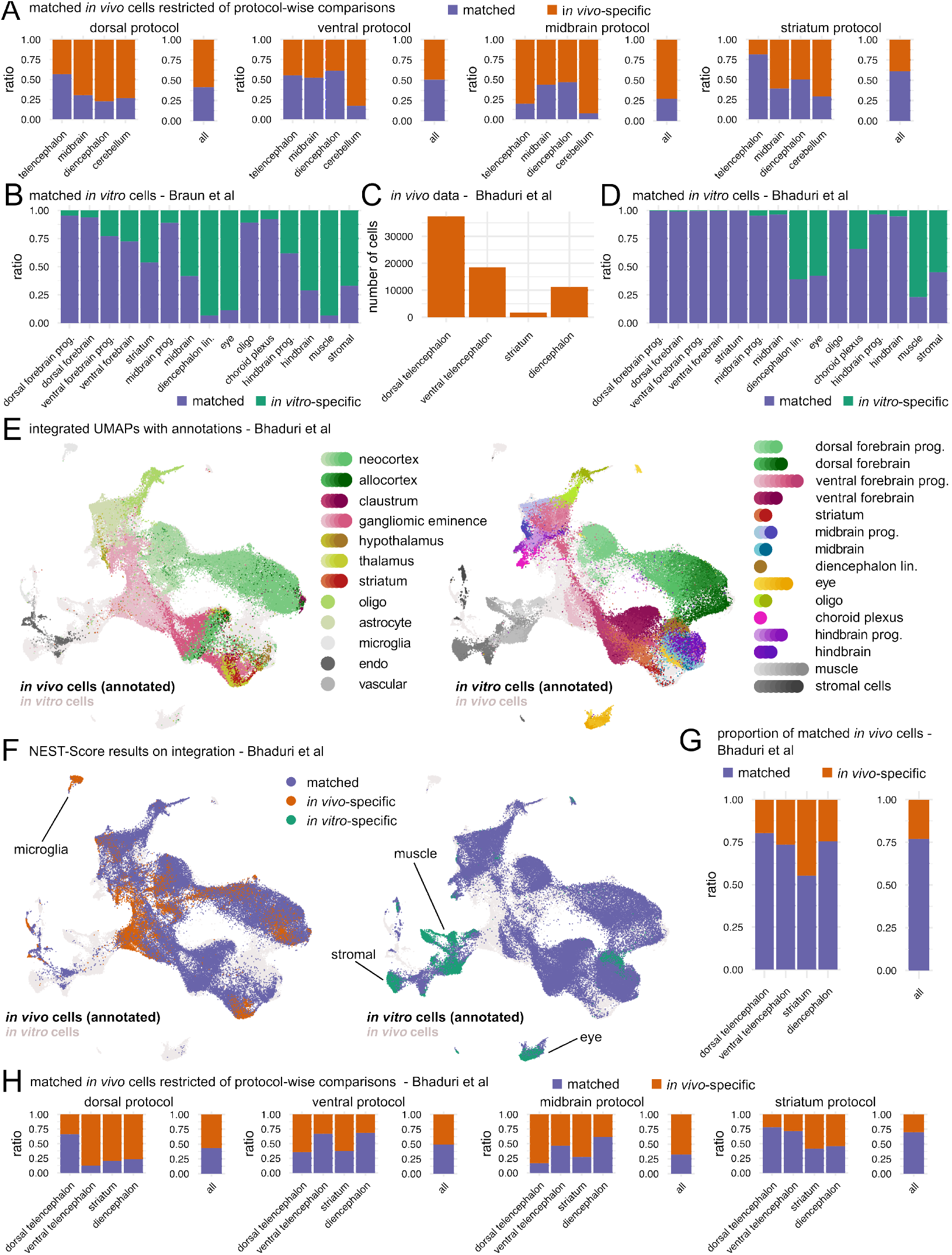
Integration of *in vivo* and *in vitro* datasets. (A) Coverage of *in vivo* cells grown in individual protocols compared to fetal brain atlas (24). (B) Coverage of well-mixed and *in vitro* specific cells per cell type after integration with *in vivo* reference data from Braun et al. (24), abbreviations: lin. = lineage, prog. = progenitors. (C) Coverage of *in vivo* cells (23) across brain regions from GW 14, 17 and 18 considered for integration. (D) As (B) but with integrated data from Bhaduri et al. (23). (E) Integrated UMAP with cell type annotation split by *in vitro* and *in vivo* (23).(F) Distribution of well-mixed, *in vivo* and *in vitro* specific cells on the integrated UMAP. (G) Ratios of *in vivo* specific and well-mixed cells per brain region. In total 77.0% of *in vivo* cells have *in vitro* cells in their local neighborhood after integration. H) Protocol-wise comparisons lead to 43.7% well-mixed cells for dorsal protocol, 49.4% for ventral protocol, 32.3% for midbrain and 70.1% for striatum protocol. (H) Coverage of *in vivo* cells grown in individual protocols compared to fetal brain atlas (23).

**Fig. S4.**
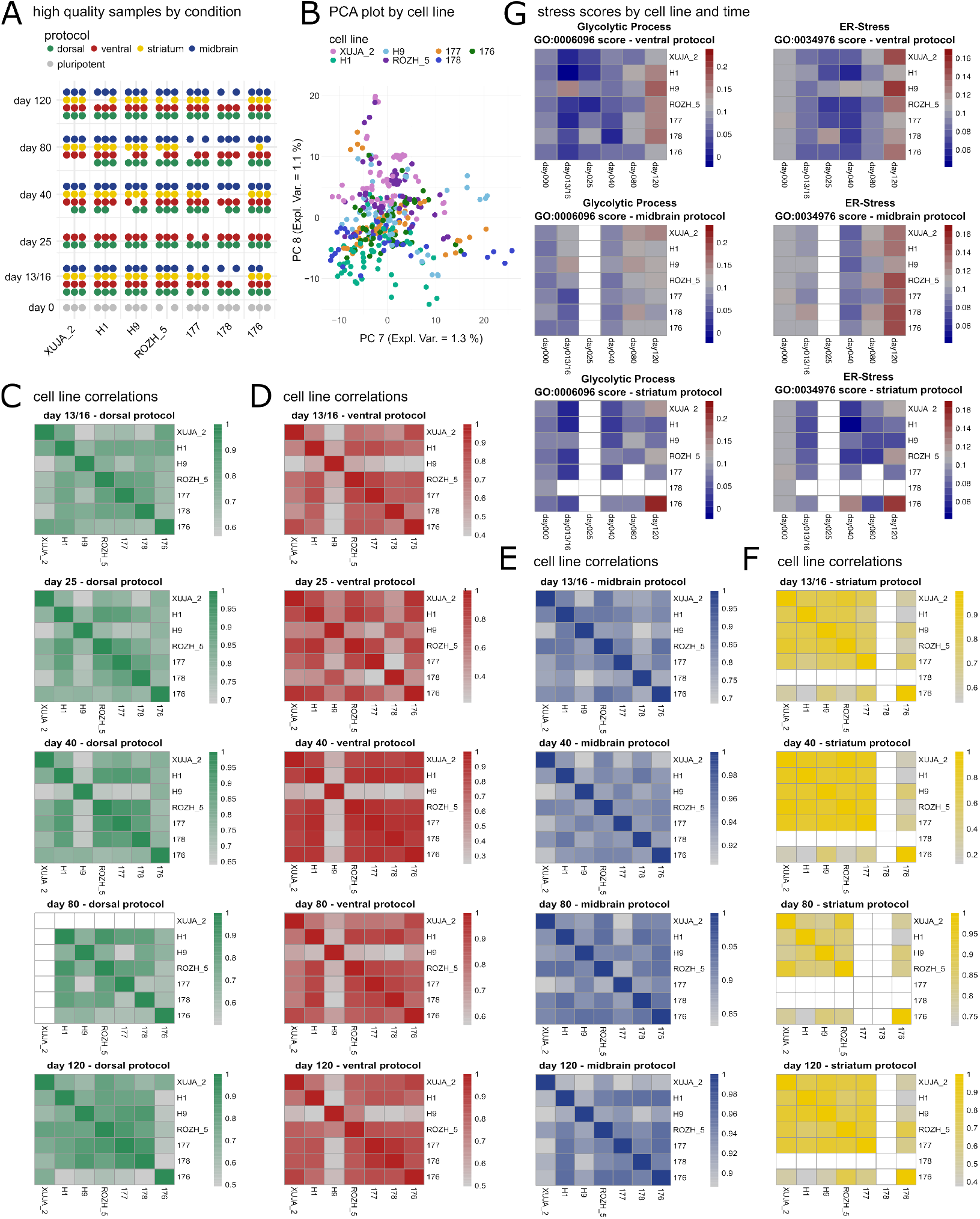
Protocol-signatures across cell lines specific, time-based bulk RNA-seq data. (A) Detailed experimental overview of bulk RNA-seq samples after quality control filtering (samples with more than 10 000 reads). (B) Principal component analysis (PCA) of bulk RNA-seq datasets. Plotted are principal components 7 and 8 with samples colored by cell line. (C) Pearson correlations between cell lines for all measured timepoints in the dorsal protocol. (D) Pearson correlations between cell lines for all measured timepoints in the ventral protocol. (E) Pearson correlations between cell lines for all measured timepoints in the midbrain protocol. (F) Pearson correlations between cell lines for all measured timepoints in the striatal protocol. (G) Average expression of GO Terms 0006096 (Glycolytic Process) and 0034976 (ER-Stress) by cell line and protocol.

**Fig. S5.**
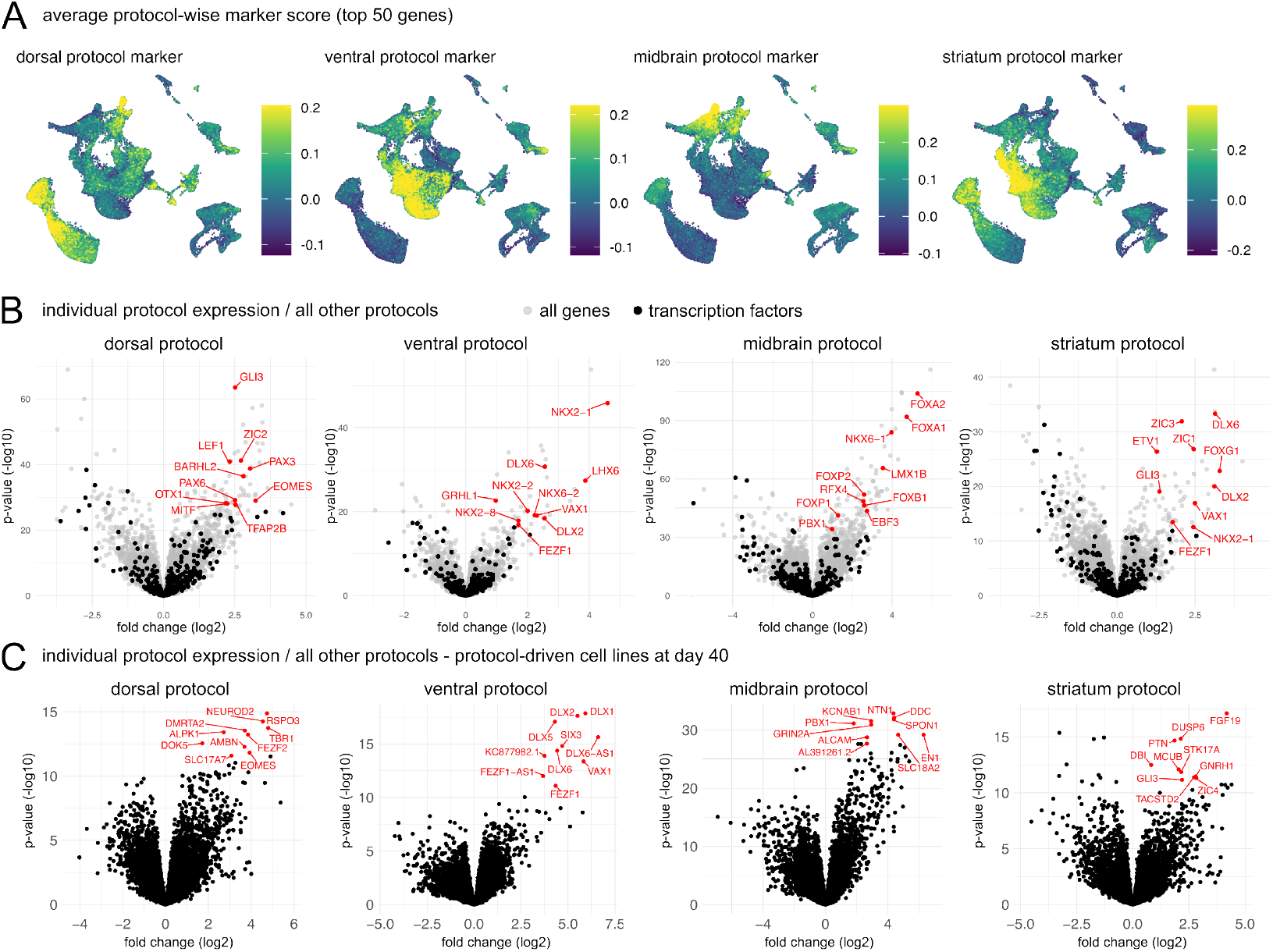
RNA-seq-based protocol signatures across protocols and early timepoint. (A) Average protocol marker scores (across all timepoints, top 50 ordered by adjusted p-value) on scRNA-seq UMAP across protocols. (B) Volcano plots of differentially expressed genes tested against all remaining three protocols separated by indicated protocol. Top 10 upregulated transcription factors are indicated. (C) Volcano plots with samples restricted to day 40 and protocol-driven cell lines within a protocol against remaining cell lines of the same protocol and all cell lines in all other protocols.

## References

1. Paola Arlotta and Fred H. Gage. Neural Organoids and the Quest to Understand and Treat Psychiatric Disease. Biological Psychiatry, 93(7):588–589, April 2023. ISSN 1873-2402. doi: 10.1016/j.biopsych.2023.01.021.

2. Paola Arlotta and Sergiu P. Paşca. Cell diversity in the human cerebral cortex: from the embryo to brain organoids. Current Opinion in Neurobiology, 56:194–198, June 2019. ISSN 1873-6882. doi: 10.1016/j.conb.2019.03.001.

3. J. Gray Camp, Farhath Badsha, Marta Florio, Sabina Kanton, Tobias Gerber, Michaela Wilsch-Bräuninger, Eric Lewitus, Alex Sykes, Wulf Hevers, Madeline Lancaster, Juergen A. Knoblich, Robert Lachmann, Svante Pääbo, Wieland B. Huttner, and Barbara Treutlein. Human cerebral organoids recapitulate gene expression programs of fetal neocortex development. Proceedings of the National Academy of Sciences, 112(51):15672–15677, December 2015. doi: 10.1073/pnas.1520760112. Publisher: Proceedings of the National Academy of Sciences.

4. Oliver L. Eichmüller and Juergen A. Knoblich. Human cerebral organoids - a new tool for clinical neurology research. Nature Reviews. Neurology, 18(11):661–680, November 2022. ISSN 1759-4766. doi: 10.1038/s41582-022-00723-9.

5. Mototsugu Eiraku, Kiichi Watanabe, Mami Matsuo-Takasaki, Masako Kawada, Shigenobu Yonemura, Michiru Matsumura, Takafumi Wataya, Ayaka Nishiyama, Keiko Muguruma, and Yoshiki Sasai. Self-Organized Formation of Polarized Cortical Tissues from ESCs and Its Active Manipulation by Extrinsic Signals. Cell Stem Cell, 3(5):519–532, November 2008. ISSN 1934-5909, 1875-9777. doi: 10.1016/j.stem.2008.09.002. Publisher: Elsevier.

6. Kevin W. Kelley and Sergiu P. Pas, ca. Human brain organogenesis: Toward a cellular understanding of development and disease. Cell, 185(1):42–61, January 2022. ISSN 1097-4172. doi: 10.1016/j.cell.2021.10.003.

7. Christina Kyrousi and Silvia Cappello. Using brain organoids to study hu-man neurodevelopment, evolution and disease. WIREs Developmental Biol - https://onlinelibrary.wiley.com/doi/pdf/10.1002/wdev.347.

8. Madeline A. Lancaster, Magdalena Renner, Carol-Anne Martin, Daniel Wenzel, Louise S. Bicknell, Matthew E. Hurles, Tessa Homfray, Josef M. Penninger, Andrew P. Jackson, and Juergen A. Knoblich. Cerebral organoids model human brain development and microcephaly. Nature, 501(7467):373–379, September 2013. ISSN 1476-4687. doi: 10.1038/nature12517. Publisher: Nature Publishing Group.

9. Madeline A. Lancaster and Juergen A. Knoblich. Organogenesis in a dish: Modeling development and disease using organoid technologies. Science, 345(6194):1247125, July 2014. doi: 10.1126/science.1247125. Publisher: American Association for the Advancement of Science.

10. Giorgia Quadrato, Juliana Brown, and Paola Arlotta. The promises and challenges of human brain organoids as models of neuropsychiatric disease. Nature Medicine, 22(11): 1220–1228, November 2016. ISSN 1546-170X. doi: 10.1038/nm.4214. Publisher: Nature Publishing Group.

11. Giorgia Quadrato and Paola Arlotta. Present and future of modeling human brain development in 3D organoids. Current Opinion in Cell Biology, 49:47–52, December 2017. ISSN 0955-0674. doi: 10.1016/j.ceb.2017.11.010.

12. Giuliana Rossi, Andrea Manfrin, and Matthias P. Lutolf. Progress and potential in organoid research. Nature Reviews. Genetics, 19(11):671–687, November 2018. ISSN 1471-0064. doi: 10.1038/s41576-018-0051-9.

13. Silvia Velasco, Amanda J. Kedaigle, Sean K. Simmons, Allison Nash, Marina Rocha, Giorgia Quadrato, Bruna Paulsen, Lan Nguyen, Xian Adiconis, Aviv Regev, Joshua Z. Levin, and Paola Arlotta. Individual brain organoids reproducibly form cell diversity of the human cerebral cortex. Nature, 570(7762):523–527, June 2019. ISSN 1476-4687. doi: 10.1038/s41586-019-1289-x.

14. Neal D. Amin, Kevin W. Kelley, Jin Hao, Yuki Miura, Genta Narazaki, Tommy Li, Patrick McQueen, Shravanti Kulkarni, Sergey Pavlov, and Sergiu P. Paşca. Generating human neural diversity with a multiplexed morphogen screen in organoids, June 2023. Pages: 2023.05.31.541819 Section: New Results.

15. Joshua A. Bagley, Daniel Reumann, Shan Bian, Julie Lévi-Strauss, and Juergen A. Knoblich. Fused cerebral organoids model interactions between brain regions. Nature Methods, 14(7):743–751, July 2017. ISSN 1548-7105. doi: 10.1038/nmeth.4304.

16. Fikri Birey, Jimena Andersen, Christopher D. Makinson, Saiful Islam, Wu Wei, Nina Huber, H. Christina Fan, Kimberly R. Cordes Metzler, Georgia Panagiotakos, Nicholas Thom, Nancy A. O’Rourke, Lars M. Steinmetz, Jonathan A. Bernstein, Joachim Hallmayer, John R. Huguenard, and Sergiu P. Paşca. Assembly of functionally integrated human forebrain spheroids. Nature, 545(7652):54–59, May 2017. ISSN 1476-4687. doi: 10.1038/nature22330. Publisher: Nature Publishing Group.

17. Christopher N. Mayhew and Richa Singhania. A review of protocols for brain organoids and applications for disease modeling. STAR Protocols, 4(1):101860, December 2022. ISSN 2666-1667. doi: 10.1016/j.xpro.2022.101860.

18. Sergiu P. Paşca. Assembling human brain organoids. Science, 363(6423):126–127, January 2019. doi: 10.1126/science.aau5729. Publisher: American Association for the Advancement of Science.

19. Daniel Reumann, Christian Krauditsch, Maria Novatchkova, Edoardo Sozzi, Sakurako Nagumo Wong, Michael Zabolocki, Marthe Priouret, Balint Doleschall, Kaja I. Ritzau-Reid, Marielle Piber, Ilaria Morassut, Charles Fieseler, Alessandro Fiorenzano, Molly M. Stevens, Manuel Zimmer, Cedric Bardy, Malin Parmar, and Jürgen A. Knoblich. In vitro modeling of the human dopaminergic system using spatially arranged ventral mid-brain–striatum–cortex assembloids. Nature Methods, 20(12):2034–2047, December 2023. ISSN 1548-7105. doi: 10.1038/s41592-023-02080-x. Publisher: Nature Publishing Group.

20. Zhisong He, Leander Dony, Jonas Simon Fleck, Artur Szałata, Katelyn X. Li, Irena Slišković, Hsiu-Chuan Lin, Malgorzata Santel, Alexander Atamian, Giorgia Quadrato, Jieran Sun, Sergiu P. Paşca, J. Gray Camp, Fabian Theis, and Barbara Treutlein. An integrated transcriptomic cell atlas of human neural organoids, October 2023. Pages: 2023.10.05.561097 Section: New Results.

21. Sabina Kanton, Michael James Boyle, Zhisong He, Malgorzata Santel, Anne Weigert, Fátima Sanchís-Calleja, Patricia Guijarro, Leila Sidow, Jonas Simon Fleck, Dingding Han, Zhengzong Qian, Michael Heide, Wieland B. Huttner, Philipp Khaitovich, Svante Pääbo, Barbara Treutlein, and J. Gray Camp. Organoid single-cell genomic atlas uncovers human-specific features of brain development. Nature, 574(7778):418–422, October 2019. ISSN 1476-4687. doi: 10.1038/s41586-019-1654-9. Publisher: Nature Publishing Group.

22. Fátima Sanchís-Calleja, Akanksha Jain, Zhisong He, Ryoko Okamoto, Charlotte Rusimbi, Pedro Rifes, Gaurav Singh Rathore, Malgorzata Santel, Jasper Janssens, Makiko Seimiya, Jonas Simon Fleck, Agnete Kirkeby, J. Gray Camp, and Barbara Treutlein. Decoding mor-phogen patterning of human neural organoids with a multiplexed single-cell transcriptomic screen, February 2024. Pages: 2024.02.08.579413 Section: New Results.

23. Aparna Bhaduri, Carmen Sandoval-Espinosa, Marcos Otero-Garcia, Irene Oh, Raymund Yin, Ugomma C. Eze, Tomasz J. Nowakowski, and Arnold R. Kriegstein. An atlas of cortical arealization identifies dynamic molecular signatures. Nature, 598(7879):200–204, Octo-ber 2021. ISSN 1476-4687. doi: 10.1038/s41586-021-03910-8. Number: 7879 Publisher: Nature Publishing Group.

24. Emelie Braun, Miri Danan-Gotthold, Lars E. Borm, Elin Vinsland, Ka Wai Lee, Peter Lön-nerberg, Lijuan Hu, Xiaofei Li, Xiaoling He, Žaneta Andrusivová, Joakim Lundeberg, Ernest Arenas, Roger A. Barker, Erik Sundström, and Sten Linnarsson. Comprehensive cell atlas of the first-trimester developing human brain, October 2022. Pages: 2022.10.24.513487 Section: New Results.

25. Jonas Simon Fleck, Fátima Sanchís-Calleja, Zhisong He, Malgorzata Santel, Michael James Boyle, J. Gray Camp, and Barbara Treutlein. Resolving organoid brain region identities by mapping single-cell genomic data to reference atlases. Cell Stem Cell, 28 (6):1148–1159.e8, June 2021. ISSN 1934-5909. doi: 10.1016/j.stem.2021.02.015.

26. Carol L. Thompson, Lydia Ng, Vilas Menon, Salvador Martinez, Chang-Kyu Lee, Katie Glattfelder, Susan M. Sunkin, Alex Henry, Christopher Lau, Chinh Dang, Raquel Garcia-Lopez, Almudena Martinez-Ferre, Ana Pombero, John L. R. Rubenstein, Wayne B. Wakeman, John Hohmann, Nick Dee, Andrew J. Sodt, Rob Young, Kimberly Smith, Thuc-Nghi Nguyen, Jolene Kidney, Leonard Kuan, Andreas Jeromin, Ajamete Kaykas, Jeremy Miller, Damon Page, Geri Orta, Amy Bernard, Zackery Riley, Simon Smith, Paul Wohnoutka, Michael J. Hawrylycz, Luis Puelles, and Allan R. Jones. A High-Resolution Spatiotemporal Atlas of Gene Expression of the Developing Mouse Brain. Neuron, 83(2):309–323, July 2014. ISSN 0896-6273. doi: 10.1016/j.neuron.2014.05.033. Publisher: Elsevier.

27. Oliver L. Eichmüller, Nina S. Corsini, Ábel Vértesy, Ilaria Morassut, Theresa Scholl, Victoria-Elisabeth Gruber, Angela M. Peer, Julia Chu, Maria Novatchkova, Johannes A. Hainfellner, Mercedes F. Paredes, Martha Feucht, and Jürgen A. Knoblich. Amplification of human interneuron progenitors promotes brain tumors and neurological defects. Science (New York, N.Y.), 375(6579):eabf5546, January 2022. ISSN 1095-9203. doi: 10.1126/science.abf5546.

28. Catarina Martins-Costa, Vincent A Pham, Jaydeep Sidhaye, Maria Novatchkova, Andrea Wiegers, Angela Peer, Paul Möseneder, Nina S Corsini, and Jürgen A Knoblich. Morpho-genesis and development of human telencephalic organoids in the absence and presence of exogenous extracellular matrix. The EMBO Journal, 42(22):e113213, November 2023. ISSN 0261-4189. doi: 10.15252/embj.2022113213. Publisher: John Wiley & Sons, Ltd.

29. Ana Uzquiano, Amanda J. Kedaigle, Martina Pigoni, Bruna Paulsen, Xian Adiconis, Kwanho Kim, Tyler Faits, Surya Nagaraja, Noelia Antón-Bolaños, Chiara Gerhardinger, Ashley Tucewicz, Evan Murray, Xin Jin, Jason Buenrostro, Fei Chen, Silvia Velasco, Aviv Regev, Joshua Z. Levin, and Paola Arlotta. Proper acquisition of cell class identity in organoids allows definition of fate specification programs of the human cerebral cortex. Cell, 185(20): 3770–3788.e27, September 2022. ISSN 1097-4172. doi: 10.1016/j.cell.2022.09.010.

30. Lukas Heumos, Anna C. Schaar, Christopher Lance, Anastasia Litinetskaya, Felix Drost, Luke Zappia, Malte D. Lücken, Daniel C. Strobl, Juan Henao, Fabiola Curion, Herbert B. Schiller, and Fabian J. Theis. Best practices for single-cell analysis across modalities. Nature Reviews Genetics, 24(8):550–572, August 2023. ISSN 1471-0064. doi: 10.1038/s41576-023-00586-w. Number: 8 Publisher: Nature Publishing Group.

31. S. Kullback and R. A. Leibler. On Information and Sufficiency. The Annals of Mathematical Statistics, 22(1):79–86, 1951. ISSN 0003-4851. Publisher: Institute of Mathematical Statistics.

32. Stefano L. Giandomenico, Magdalena Sutcliffe, and Madeline A. Lancaster. Generation and long-term culture of advanced cerebral organoids for studying later stages of neural development. Nature Protocols, 16(2):579–602, February 2021. ISSN 1750-2799. doi: 10.1038/s41596-020-00433-w. Publisher: Nature Publishing Group.

33. Tim Stuart, Andrew Butler, Paul Hoffman, Christoph Hafemeister, Efthymia Papalexi, William M. Mauck, Yuhan Hao, Marlon Stoeckius, Peter Smibert, and Rahul Satija. Comprehensive Integration of Single-Cell Data. Cell, 177(7):1888–1902.e21, June 2019. ISSN 0092-8674. doi: 10.1016/j.cell.2019.05.031.

34. Gabriel E. Hoffman and Eric E. Schadt. variancePartition: interpreting drivers of variation in Naas, and Balmãna et al. | Reconstitution of Human Brain Cell Diversity in Organoids via Four Protocols complex gene expression studies. BMC Bioinformatics, 17(1):1–13, December 2016. ISSN 1471-2105. doi: 10.1186/s12859-016-1323-z. Number: 1 Publisher: BioMed Central.

35. Matthew E. Ritchie, Belinda Phipson, D. Wu, Yifang Hu, Charity W. Law, Wei Shi, and Gordon K. Smyth. imma powers differential expression analyses for RNA-sequencing and microarray studies. Nucleic Acids Research, 43(7):e47, April 2015. ISSN 0305-1048. doi: 10.1093/nar/gkv007.

36. Aparna Bhaduri, Madeline G. Andrews, Walter Mancia Leon, Diane Jung, David Shin, Denise Allen, Dana Jung, Galina Schmunk, Maximilian Haeussler, Jahan Salma, Alex A. Pollen, Tomasz J. Nowakowski, and Arnold R. Kriegstein. Cell stress in cortical organoids impairs molecular subtype specification. Nature, 578(7793):142–148, February 2020. ISSN 1476-4687. doi: 10.1038/s41586-020-1962-0. Publisher: Nature Publishing Group.

37. Aaron Gordon, Se-Jin Yoon, Stephen S. Tran, Christopher D. Makinson, Jin Young Park, Jimena Andersen, Alfredo M. Valencia, Steve Horvath, Xinshu Xiao, John R. Huguenard, Sergiu P. Pas, ca, and Daniel H. Geschwind. Long-term maturation of human cortical organoids matches key early postnatal transitions. Nature Neuroscience, 24(3):331–342, March 2021. ISSN 1546-1726. doi: 10.1038/s41593-021-00802-y. Publisher: Nature Publishing Group.

38. Xuyu Qian, Hongjun Song, and Guo-li Ming. Brain organoids: advances, applications and challenges. Development, 146(8):dev166074, April 2019. ISSN 0950-1991. doi: 10.1242/dev.166074.

39. Ábel Vértesy, Oliver L Eichmüller, Julia Naas, Maria Novatchkova, Christopher Esk, Meritxell Balmaña, Sabrina Ladstaetter, Christoph Bock, Arndt von Haeseler, and Juergen A Knoblich. Gruffi: an algorithm for computational removal of stressed cells from brain organoid transcriptomic datasets. The EMBO Journal, 41(17):e111118, September 2022. ISSN 0261-4189. doi: 10.15252/embj.2022111118. Publisher: John Wiley & Sons, Ltd.

40. Christopher Esk, Dominik Lindenhofer, Simon Haendeler, Roelof A. Wester, Florian Pflug, Benoit Schroeder, Joshua A. Bagley, Ulrich Elling, Johannes Zuber, Arndt von Haeseler, and Jürgen A. Knoblich. A human tissue screen identifies a regulator of ER secretion as a brain-size determinant. Science, 370(6519):935–941, November 2020. doi: 10.1126/science.abb5390. Publisher: American Association for the Advancement of Science.

41. Jonas Simon Fleck, Sophie Martina Johanna Jansen, Damian Wollny, Fides Zenk, Makiko Seimiya, Akanksha Jain, Ryoko Okamoto, Malgorzata Santel, Zhisong He, J. Gray Camp, and Barbara Treutlein. Inferring and perturbing cell fate regulomes in human brain organoids. Nature, 621(7978):365–372, September 2023. ISSN 1476-4687. doi: 10.1038/s41586-022-05279-8. Publisher: Nature Publishing Group.

42. Stefania Fedele, Ginetta Collo, Katharina Behr, Josef Bischofberger, Stephan Müller, Tilo Kunath, Klaus Christensen, Anna Lisa Gündner, Martin Graf, Ravi Jagasia, and Verdon Taylor. Expansion of human midbrain floor plate progenitors from induced pluripotent stem cells increases dopaminergic neuron differentiation potential. Scientific Reports, 7(1):6036, July 2017. ISSN 2045-2322. doi: 10.1038/s41598-017-05633-1. Publisher: Nature Publishing Group.

43. Winston Chang, Joe Cheng, JJ Allaire, Carson Sievert, Barret Schloerke, Yihui Xie, Jeff Allen, Jonathan McPherson, Alan Dipert, and Barbara Borges. shiny: Web Application Framework for R, 2024.

44. Se-Jin Yoon, Lubayna S. Elahi, Anca M. Pas, ca, Rebecca M. Marton, Aaron Gordon, Omer Revah, Yuki Miura, Elisabeth M. Walczak, Gwendolyn M. Holdgate, H. Christina Fan, John R. Huguenard, Daniel H. Geschwind, and Sergiu P. Pas, ca. Reliability of human cortical organoid generation. Nature Methods, 16(1):75–78, January 2019. ISSN 1548-7105. doi: 10.1038/s41592-018-0255-0. Publisher: Nature Publishing Group.

45. Maren Büttner, Zhichao Miao, F. Alexander Wolf, Sarah A. Teichmann, and Fabian J. Theis. A test metric for assessing single-cell RNA-seq batch correction. Nature Methods, 16(1): 43–49, January 2019. ISSN 1548-7105. doi: 10.1038/s41592-018-0254-1. Number: 1 Publisher: Nature Publishing Group.

46. Reiner A. Wimmer, Alexandra Leopoldi, Martin Aichinger, Nikolaus Wick, Brigitte Hantusch, Maria Novatchkova, Jasmin Taubenschmid, Monika Hämmerle, Christopher Esk, Joshua A. Bagley, Dominik Lindenhofer, Guibin Chen, Manfred Boehm, Chukwuma A. Agu, Fengtang Yang, Beiyuan Fu, Johannes Zuber, Juergen A. Knoblich, Dontscho Kerjaschki, and Josef M. Penninger. Human blood vessel organoids as a model of diabetic vasculopathy. Nature, 565(7740):505–510, January 2019. ISSN 1476-4687. doi: 10.1038/s41586-018-0858-8.

47. Alexander Atamian, Marcella Birtele, Negar Hosseini, Tuan Nguyen, Anoothi Seth, Ashley Del Dosso, Sandeep Paul, Neil Tedeschi, Ryan Taylor, Marcelo P. Coba, Ranmal Samarasinghe, Carlos Lois, and Giorgia Quadrato. Human cerebellar organoids with functional Purkinje cells. Cell Stem Cell, 31(1):39–51.e6, January 2024. ISSN 1875-9777. doi: 10.1016/j.stem.2023.11.013.

48. Haynes Heaton, Arthur M. Talman, Andrew Knights, Maria Imaz, Daniel J. Gaffney, Richard Durbin, Martin Hemberg, and Mara K. N. Lawniczak. Souporcell: robust clustering of single-cell RNA-seq data by genotype without reference genotypes. Nature Methods, 17(6):615–620, June 2020. ISSN 1548-7105. doi: 10.1038/s41592-020-0820-1. Publisher: Nature Publishing Group.

49. Yuhan Hao, Tim Stuart, Madeline H. Kowalski, Saket Choudhary, Paul Hoffman, Austin Hartman, Avi Srivastava, Gesmira Molla, Shaista Madad, Carlos Fernandez-Granda, and Rahul Satija. Dictionary learning for integrative, multimodal and scalable single-cell analysis. Nature Biotechnology, 42(2):293–304, February 2024. ISSN 1546-1696. doi: 10.1038/s41587-023-01767-y. Publisher: Nature Publishing Group.

50. Itay Tirosh, Benjamin Izar, Sanjay M. Prakadan, Marc H. Wadsworth, Daniel Treacy, John J. Trombetta, Asaf Rotem, Christopher Rodman, Christine Lian, George Murphy, Mohammad Fallahi-Sichani, Ken Dutton-Regester, Jia-Ren Lin, Ofir Cohen, Parin Shah, Diana Lu, Alex S. Genshaft, Travis K. Hughes, Carly G. K. Ziegler, Samuel W. Kazer, Aleth Gaillard, Kellie E. Kolb, Alexandra-Chloé Villani, Cory M. Johannessen, Aleksandr Y. Andreev, Eliezer M. Van Allen, Monica Bertagnolli, Peter K. Sorger, Ryan J. Sullivan, Keith T. Flaherty, Dennie T. Frederick, Judit Jané-Valbuena, Charles H. Yoon, Orit Rozenblatt-Rosen, Alex K. Shalek, Aviv Regev, and Levi A. Garraway. Dissecting the multicellular ecosystem of metastatic melanoma by single-cell RNA-seq. Science, 352(6282):189–196, April 2016. doi: 10.1126/science.aad0501. Publisher: American Association for the Advancement of Science.

